# Fast actin disassembly and fimbrin mechanosensitivity support rapid turnover during clathrin-mediated endocytosis

**DOI:** 10.1101/2022.11.25.517735

**Authors:** Sayed Iman Mousavi, Michael M. Lacy, Xiaobai Li, Julien Berro

## Abstract

The actin cytoskeleton is central to force production in numerous cellular processes in eukaryotic cells. During clathrin-mediated endocytosis (CME), a dynamic actin meshwork is required to deform the membrane against high membrane tension or turgor pressure. Previous experimental work from our lab showed that several endocytic proteins, including actin and actin-interacting proteins, turn over several times during the formation of a vesicle during CME in yeast, and their dwell time distributions were reminiscent of Gamma distributions with a peak around 1 s (Lacy et al., 2019). However, the distribution for the filament crosslinking protein fimbrin contains a second peak around 0.5 s. To better understand the nature of these dwell time distributions, we developed a stochastic model for the dynamics of actin and its binding partners. Our model demonstrates that very fast actin filament disassembly is necessary to reproduce experimental dwell time distributions. Our model also predicts that actin-binding proteins bind rapidly to nascent filaments and filaments are fully decorated. Last, our model predicts that fimbrin detachment from actin endocytic structures is mechanosensitive to explain the extra peak observed in the dwell time distribution.

## Introduction

Eukaryotic cells rely on clathrin-mediated endocytosis (CME) to produce membrane-bound vesicles that transport materials from the cell surface into the cytoplasm. Dozens of proteins rapidly self-assemble and coordinate to reshape the plasma membrane into a vesicle (Goode et al., 2015; Kaksonen & Roux, 2018; Lacy et al., 2018). In yeast, a localized, dynamic actin meshwork is required to generate forces to counteract high turgor pressure (Aghamohammadzadeh & Ayscough, 2009; Boulant et al., 2011; Collins et al., 2011). Despite numerous experimental and theoretical studies to characterize the role of endocytic proteins, our understanding of the overall mechanics of endocytosis remains incomplete (Lacy et al., 2018). Quantitative fluorescence microscopy studies of CME in fission and budding yeasts have characterized the robust recruitment timelines of coat proteins, actin, actin-associated proteins, and membrane scission factors (Berro & Pollard, 2014a; Kaksonen & Roux, 2018; Mund et al., 2018; Picco et al., 2015; Sirotkin et al., 2010). Computational and mathematical modeling has enabled a deeper understanding of the mechanisms of CME, demonstrating that the dendritic nucleation model for actin dynamics can recapitulate the observed dynamics of assembly and disassembly (Berro et al., 2010), and proposing mechanisms to explain how the actin meshwork generates, stores, and transmits force to deform the plasma membrane (Carlsson, 2018; Carlsson & Bayly, 2014; Dmitrieff & Nédélec, 2016; Ma & Berro, 2018, 2019; Nickaeen et al., 2019). Yet, many molecular details for the dynamics of force production during CME remain unclear.

Using single-molecule microscopy in fission yeast, we recently revealed the rapid and continuous exchange of endocytic molecules at sites of endocytosis (Lacy et al., 2019), indicating that the actin meshwork and membrane coat proteins turn over multiple times during a single endocytic event. Whereas endocytic actin patches have a lifetime of around 20 seconds, the dwell times of individual molecules were 1 to 2 seconds on average (Lacy et al., 2019). The single-molecule dwell time distributions followed a rise and decay reminiscent of a Gamma distribution, with small but statistically significant differences between some proteins. Since the single-molecule dwell time distributions for most actin-associated proteins look close to each other, we hypothesized that these dynamics were dictated by the underlying kinetics of the actin subunits to which they are bound. Interestingly, the dwell time distribution for the actin filament crosslinking protein fimbrin contained an additional peak of short-lived events (hereafter called the “fast peak”; the other peak, which is common to all distributions will be referred to as the “slow peak”). Such rapid turnover of the endocytic machinery components has important implications for the biophysical mechanisms of CME, but the experimental data are difficult to interpret directly due to the convoluted kinetics and numerous components in the endocytic structures.

To identify models consistent with the dwell time distributions observed in Lacy et al., (2019), we carried out computational simulations of actin filaments containing various biochemical components under different hypotheses. We demonstrate that fast actin disassembly is the reason for fast turnover, and the detachment of filaments from the actin network plays a minimal role. Using values for the rate constants of actin filament depolymerization and severing that recapitulate the observed dwell time distributions of actin monomers, we show that the dynamics of the actin-bound proteins are the result of the dynamics of the actin subunit it is bound to. We demonstrate that the fast peak in the experimental dwell time distribution of fimbrin is not caused by a sub-population of fimbrin molecules that would participate in cytokinesis, actin cables, or other actin structures. Rather, this fast peak can be explained by a force-dependent unbinding of fimbrin at CME sites. Overall, our results contribute important insights into the biomechanical and biochemical properties of the endocytic actin meshwork and provide a new context for ongoing experimental and theoretical work in endocytosis and other actin-based structures.

## Results

### Actin polymerization and depolymerization with standard rate constants alone cannot explain the peaked dwell time distribution of actin subunits at CME sites

Our previous single-molecule experiments demonstrated that the dwell times for actin molecules at sites of endocytosis follow a peaked distribution reminiscent of a Gamma distribution with an average of around 1.5 s (Lacy et al., 2019). We speculated these distributions were the result of the complex dynamics of polymerization, depolymerization, filament severing, and turnover. However, precisely interpreting the experimental measurements of these single-molecule dwell times and testing hypothetical mechanisms require mathematical modeling. In the commonly accepted organization of the endocytic actin meshwork, filaments are nucleated at the base of the clathrin-coated pit by the Arp2/3 complex and the overall actin structure grows essentially parallel to the axis of invagination (Mund et al., 2018; Nickaeen et al., 2019). In this study, we assume there are no significant spatial differences in the biochemical properties of actin filaments within the meshwork, as no experimental data suggesting otherwise has been reported. Therefore, it is not necessary to simulate the entire meshwork as we can approximate the dwell time distributions from simulations of individual filaments.

In this study, we developed stochastic simulations of actin filaments, initially only including polymerization, depolymerization, and aging of the bound nucleotide (**Fig. 1**A), and later with increasing complexity to test new hypotheses. We used the rejection sampling algorithm (Vestergaard & Génois, 2015) for all the reactions. We also assumed that only one percent of the proteins were tagged to match the experimental conditions of the data we used for comparison (Lacy et al., 2019). Note that this assumption also minimizes the correlations between the simulated data points that would happen otherwise. The dwell time of an individual actin subunit was determined as the time interval it spent as part of a filament from the time it is polymerized to the time it dissociates from the filament, either by depolymerization or after a filament fragment containing the subunit has been dissociated from the actin-network.

**Figure 1.**
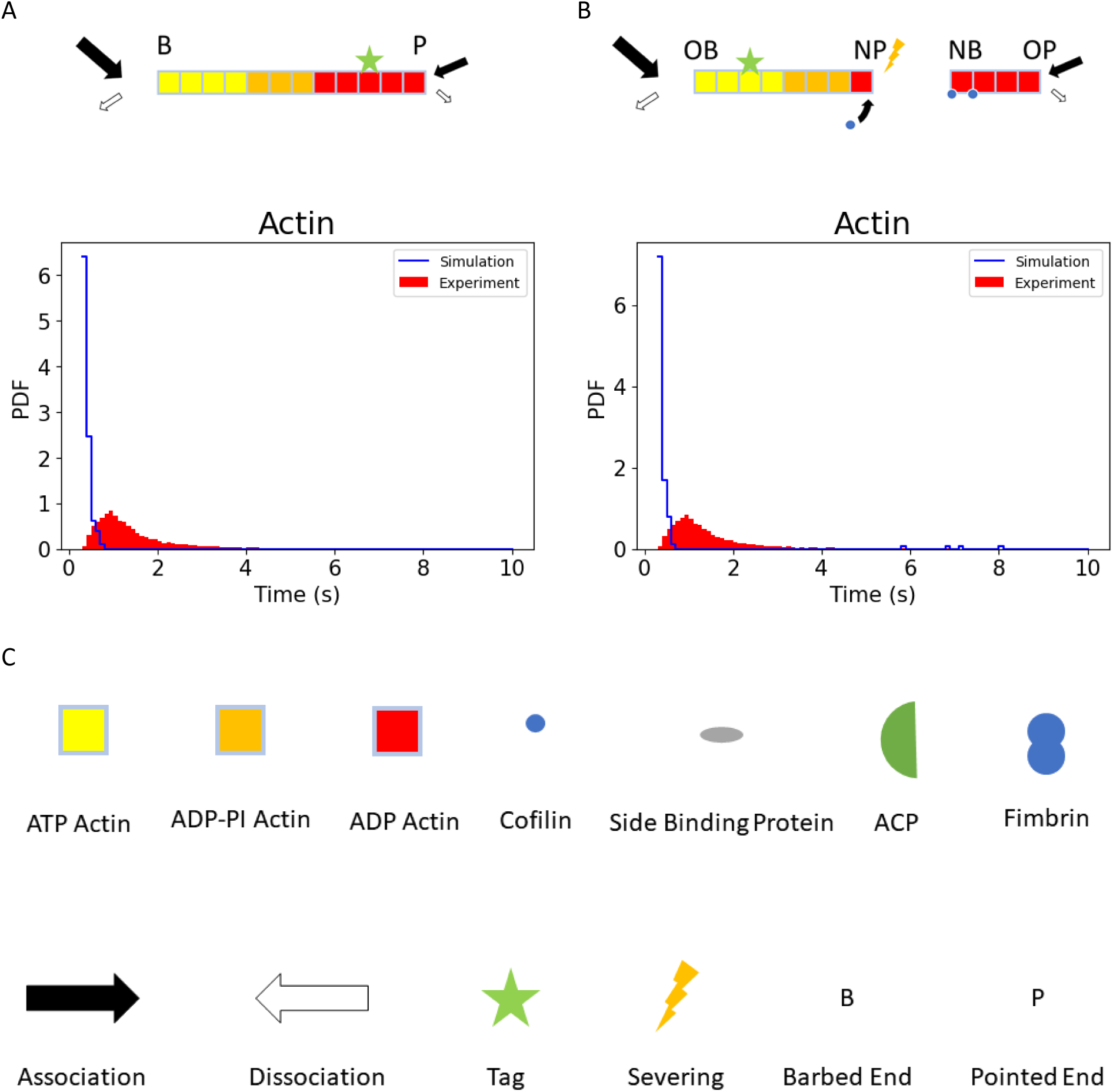
Actin dynamics with standard rate constants cannot produce peaked actin subunit dwell time distributions. A and B) The dwell time of actin subunits within a filament depends on the rate constants for polymerization, depolymerization, aging, and severing. Individual filaments were simulated using the rate constants and concentrations in **Table 2**. A) In the absence of severing, the dwell times of actin subunits follow an exponential distribution. Inset: distribution with finer bins between 0 and 1 second. B) Adding severing (e.g., by ADF/cofilin) leads to peaked actin dwell time distributions. However, the average dwell time is much larger than observed in the experimental data. There is no statistically significant difference between the simulated distributions in panels A and B as the probability of severing a filament within 10 seconds is very low when we use standard rate constants. Blue: simulated distributions; Red: experimental data for actin (Act1p) (adapted from Lacy et al., (2019). C) Symbols used in the schematics of the paper; OB/NB: Old/New Barbed end, OP/NP: Old/New -pointed end.

**Table 1.** List of reactions.

| # | Reaction | Description | Rate constants |
| --- | --- | --- | --- |
| 1 | ATP-G-Actin+Barbed_end ↔ F-actin-ATP | Polymerization and depolymerization of an ATP-actin monomer at the barbed end | $k_{Polym\_barbed\_ATP}^+$,<br>$k_{Polym\_barbed\_ATP}^-$ |
| 2 | ADP-G-Actin+ Barbed_end ↔ F-actin-ADP | Polymerization and depolymerization of an ADP-actin monomer at the barbed end | $k_{Polym\_barbed\_ADP}^+$,<br>$k_{Polym\_barbed\_ADP}^-$ |
| 3 | ATP-G-Actin+Pointed_end ↔ F-actin-ATP | Polymerization and depolymerization of an ATP-actin monomer at the pointed end | $k_{Polym\_pointed\_ATP}^+$,<br>$k_{Polym\_pointed\_ATP}^-$ |
| 4 | ADP-G-Actin+ Pointed_end ↔ F-actin-ADP | Polymerization and depolymerization of an ADP-actin monomer at the pointed end | $k_{Polym\_barbed\_ADP}^+$,<br>$k_{Polym\_barbed\_ADP}^-$ |
| 5 | F-actin-ATP → F-actin-ADP-Pi | Hydrolysis of ATP in actin subunits | $k_{ATP\ hydrolysis}$ |
| 6 | ADP-Pi subunit → ADP subunit | Pi Release | Bare filament?<br>$k_{ADP-Pi}$ :<br>$k_{ADP-Pi\ cooperative\ release}^*$ |
| 9 | Cof+Bare subunit | Add cofilin to a bare subunit | [ADF] × (the adjacent subunit is ADF/cofilin-bound)?<br>$k_{cofilin\_cooperative}^+$:<br>$k_{cofilin}^+$ |
| 10 | Cof → ∅ | Remove cofilin from a cofilinated subunit | $k_{cofilin}^-$ |
| 11 | Cofilactin filament → 2 Cofilactin filament | Sever a cofilactin filament into two | $k_{cofilin}^{severing}$ |
| 12 | ACP+F_barbed → Capped barbed end | Cap the uncap barbed end | The adjacent subunit is ADF/cofilin-bound? $\frac{k_{Cap}^+}{10}$:<br>$k_{Cap}^+$ |
| 13 | Capped barbed end → F_barbed + ACP | Uncap the capped barbed end | $k_{Cap}^-$ |
| *X==Y?value1:value2 means if the condition is satisfied, we used the value1 otherwise we use value2 |  |  |  |

**Table 2.** Reaction rate constants and concentration.

| Parameter | Optimal Value | Unit | Reference Value | Reference |
| --- | --- | --- | --- | --- |
| [ATP-G-Actin] | 10* | $\mu\text{M}$ | 21.6 | Sirotkin et al 2010 |
| [ADP-G-Actin] | 0 | $\mu\text{M}$ | | |
| [Cap] | 0.8 | $\mu\text{M}$ | 0.8 | Sirotkin et al 2010 |
| [Cofilin] | 150 | $\mu\text{M}$ | 40 | Sirotkin et al 2010 |
| $k_{Polym\_barbed\_ATP}^+$ | 11.6 | $\mu\text{M}^{-1}\text{s}^{-1}$ | 11.6 | Pollard 1986. |
| $k_{Polym\_barbed\_ADP}^+$ | 3.8 | $\mu\text{M}^{-1}\text{s}^{-1}$ | 3.8 | Pollard 1986. |
| $k_{Polym\_barbed\_ATP}^-$ | 1 | $\text{s}^{-1}$ | 1.4 | Pollard 1986. |
| $k_{Polym\_barbed\_ADP}^-$ | 10 | $\text{s}^{-1}$ | 7.2 | Pollard 1986. |
| $k_{Polym\_pointed\_ATP}^+$ | 0 | $\mu\text{M}^{-1}\text{s}^{-1}$ | 1.3 | Pollard 1986. |
| $k_{Polym\_pointed\_ADP}^+$ | 0 | $\mu\text{M}^{-1}\text{s}^{-1}$ | 0.16 | Pollard 1986. |
| $k_{Polym\_pointed\_ATP}^-$ | 10 | $\text{s}^{-1}$ | 0.8 | Pollard 1986. |
| $k_{Polym\_pointed\_ADP}^-$ | 10 | $\text{s}^{-1}$ | 0.27 | Pollard 1986. |
| $k_{cofilin}^+$ | 0.0085 | $\mu\text{M}^{-1}\text{s}^{-1}$ | 0.0085 | Blanchoin et al. (1999) |
| $k_{cofilin\_cooperative}^+$ | 0.075 | $\mu\text{M}^{-1}\text{s}^{-1}$ | 0.075 | Sirotkin et al 2010 |
| $k_{cofilin}^-$ | 0.005 | $\text{s}^{-1}$ | 0.005 | Blanchoin et al. (1999) |
| $k_{severing\_cofilin}$ | 1.24 | $\text{s}^{-1}$ | 0.003 | Sirotkin et al 2010 |
| $k_{ATP\ hydrolysis}$ | 30 | $\text{s}^{-1}$ | 0.35 | Sirotkin et al 2010 |
| $k_{ADP-PI\ release}$ | 0.2 | $\text{s}^{-1}$ | 0.0019 | Sirotkin et al 2010 |
| $k_{ADP-PI\ cooperative\ release}$ | 3 | $\text{s}^{-1}$ | 0.035 | Sirotkin et al 2010 |
| $k_{Cap}^+$ | 7 | $\mu\text{M}^{-1}\text{s}^{-1}$ | 7 | Sirotkin et al 2010 |
| $k_{Cap}^-$ | 0.004 | $\text{s}^{-1}$ | 0.004 | Kuhn and Pollard (2007) |
| $k_{Cap-cofilin-actin-barbed}^-$ | 4 | $\text{s}^{-1}$ | ~4 | Wioland et al 2017 |
| $k_{fimbrin-diffusive}^+$ | 1.4 | $s^{-1}$ | | This study |
| $k_{fimbrin-one-side-bounded}^+$ | 0.7 | $s^{-1}$ | | This study |
| $k_{other\ proteins\ competer\ fimbrin}^+$ | 20 | $s^{-1}$ | | This study |
| $k_{fimbrin-crosslinked-zero-load}^-$ | 1.8 | $s^{-1}$ | | This study |
| $k_{fimbrin-one-side-bounded}^-$ | 0.017 | $s^{-1}$ | | This study |
| $k_{other\ proteins\ competer\ fimbrin}^-$ | 0 | $s^{-1}$ | | This study |
| Potential Energy Scale Intercept | 0.082 | J |  | This study |
| Potential Energy Shape Intercept | 0.064 | J |  | This study |
| Potential Energy Scale Slope | 27.33 | J/s |  | This study |
| Potential Energy Shape Slope | 2.6 | J/s |  | This study |
| *It is estimated that the applied force on actin filaments is between 1 to 10 pN. Using $r_{on} = k_{on} [ATP - G - Actin] \exp (F\delta/K_bT)$, and substituting $\delta = 2.75\ nm$, $F = 1\ pN$, we get $r_{on} \approx (0.5 [ATP - G - Actin])k_{on}$ |  |  |  |  |

In the absence of filament severing, as filaments can polymerize and depolymerize from both ends, most sets of parameters create filaments that grow indefinitely. In these conditions, virtually all the subunits remained in the filament for the entire simulation time. In this case, only the dwell times of the rare subunits that dissociate from the filament just after they have been polymerized and before a new subunit is polymerized contribute to the dataset. The distribution of these events follows an exponential (**Fig. 1**A).

### Severing of actin filaments produces a peaked dwell time distribution

Next, we added to our simulations a filament severing process, e.g., by ADF/cofilin, a mechanism known to produce filaments of any average length, depending on rates and concentrations, and increases the rate of turnover (Michelot & Drubin, 2011; Mohapatra et al., 2016; Roland et al., 2008). We implemented ADF/cofilin binding kinetics and severing similarly to previous models (Berro et al., 2007; Michelot & Drubin, 2011; Mohapatra et al., 2016; Roland et al., 2008) (**Fig. 1**B). As in Roland et al. (2008), after severing the filament, we discarded all the subunits of the fragment that contained the pointed end before the severing event (called the “old-pointed end”). In addition to severing, we tested the hypothesis that binding of ADF/cofilin to the ends of a filament destabilizes the filament ends and halts the polymerization (Wioland et al., 2019). Using these hypotheses, we were able to obtain dwell time distributions of actin subunits that are peak-shaped with a long tail, qualitatively reminiscent of the experimental data from Lacy et al., (2019) Using published cellular concentrations and rate constants measured *in vitro* and using unrealistically long simulation times (∼1000 s), the average dwell times were markedly larger, and the peaks of the distribution were broader than observed in the experimental data (**Fig. S1**A inset). However, when we used realistic simulation times comparable to the lifetime of actin endocytic structures (10 seconds), the dwell time distributions were not notably different from the distributions obtained without severing as the probability of severing a filament in 10 seconds was very low.

Up to this point, we did not include capping proteins in our simulations. Adding barbed end capping did not significantly change the results if data were collected within ∼10 seconds of simulated time. However, after 10 seconds, the filaments are still in a transient state and have not yet reached a dynamic steady state. Within this dynamic steady state, the filaments undergo growth, severing, and shrinking, but the statistical distributions of their lengths and internal biochemical states remain constant **(Fig. S1** and **S2)**. In the conditions used in this section, the dynamic steady state was reached after ∼100 seconds of simulated time (**Fig. S2**B and D), and a peak in the dwell time distributions appeared between 20 seconds to 40 seconds, depending on the presence of capping proteins (**Fig. S1**A and B). Counterintuitively, the mean dwell time was longer when capping proteins were present in the simulations because the average filament growth rate was reduced, which consequently decreased the number of available ADF/cofilin binding sites, reducing the binding probability of ADF/cofilin and therefore the filament severing rate. After increasing the concentrations and all the rate constants of ADF/cofilin binding and severing by a factor of ten, the peak of the dwell time distribution decreased from 20 seconds to ∼10 seconds (**Fig. S1**C), which is still an order of magnitude higher than the value we measured in our experiments. To obtain these results, simulations had to run for more than ∼100 seconds before the filament lengths and the dwell time distributions reached a steady state (**Fig. S1**D). This time is longer than the time it takes for actin to fully assemble during CME (∼10 seconds). In addition, the filaments’ length was 420 ± 80 *nm* after 10 seconds of simulations (**Fig. S2**B and D), which is significantly larger than the values (∼50-100 *nm*) measured by electron microscopy (Rodal et al., 2005; Young et al., 2004) or the values (∼50-180 *nm*) estimated from quantitative microscopy and mathematical modeling (Berro et al., 2010; Berro & Pollard, 2014b). We conclude that while adding ADF/cofilin and capping proteins to the model makes the simulated dwell time distributions of actin subunits appear similar to the experimental data, the standard rate constants for polymerization, depolymerization, barbed end capping, and severing from published sources cannot produce the short dwell times of actin subunits during CME.

### Very fast filament disassembly is required to decrease the average dwell time of actin subunits

Using the depolymerization and severing rate constants measured *in vitro* (Blanchoin et al., 2000; Fujiwara et al., 2007; Pollard, 1986), the average dwell time of actin monomers was longer than the reported value in Lacy et al., (2019). One possible explanation is that in endocytic patches *in vivo*, a fraction of severed filaments detaches and rapidly moves away from the actin network, therefore decreasing the average dwell time of actin subunits (Berro et al., 2010; Chen & Pollard, 2013; Sirotkin et al., 2010). As our simulations did not account for the whole network of actin filaments, we approximated this effect using a reaction with a rate that can be time-dependent or constant (see **Materials and Methods** section). Upon triggering this reaction, we removed all the subunits of the released filaments from the simulation and added the removed tagged subunits to the dwell time distribution. To explore this possibility, we investigated two different concentrations (40 and 150 *μM*) for ADF/cofilin and three different severing rates (0.003, 0.03, and 0.3 *s*^−1^). For each of these conditions, we screened a range of time-dependent detachment rates as described in the **Materials and Methods** section. To compare the simulated and experimental distributions, we used Cohen’s *w*, which is defined as 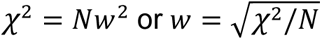, where *χ*^2^ is the chi-square between both distributions and *N* is the total number of dwell time measurements. Values of *w* around 0.1, 0.3 and 0.5 correspond, respectively, to small, medium, and large effect sizes. Using the reference values of **Table 2**, the best fit had a *w* value of 0.42 (**Fig. 2**A and **Fig. S3**A) and the minimum *w* value of all the six different configurations was 0.29 (**Fig. S3**).

**Figure 2.**
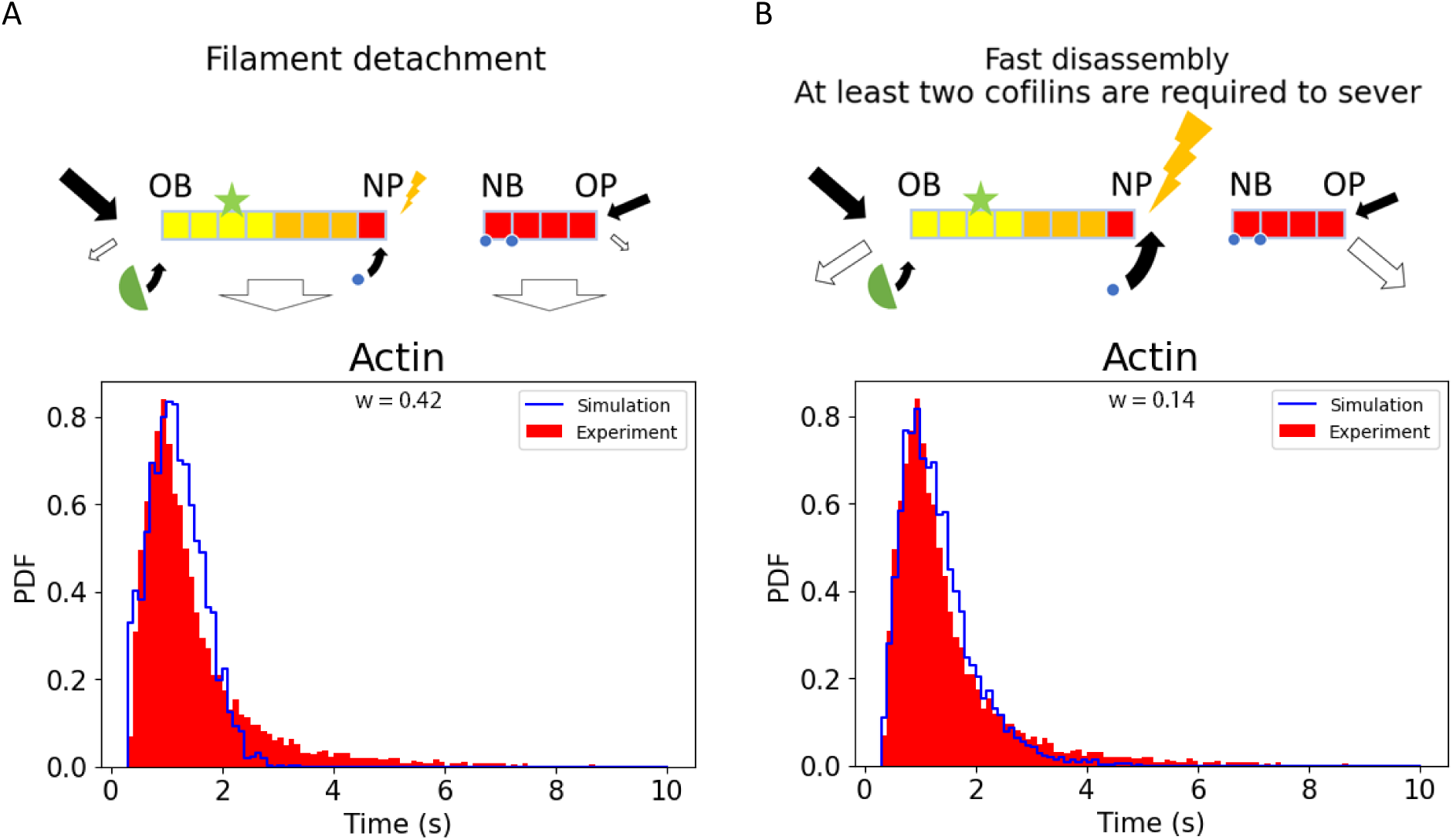
Fast removal of actin subunits is necessary to obtain peaked dwell time distributions with a small average. A) Severed filaments were randomly removed from the simulation, using *in vivo* depolymerization and severing rates. The removal of a fragment was triggered by a reaction with a time-dependent reaction rate using *filamentDetachmentRate* = *max*{0, *slop*_*detachmentRate*_ ∗ (*filamentAge*) + *intercept*_*detachmentRate*_ }, where *filamentAge* = *current time* − *mean*(*subunit polymerization time*), *slop*_*detachmentRate*_ determines the rate of *filamentDetachmentRate* changes, and *intercept*_*detachmentRate*_ with positive values determines the *filamentDetachmentRate* right after a new filament emerges in the actin-network, while negative values determine the time that the *filamentDetachmentRate* remains zero after a new actin filament is formed. The best fit (*w* = 0.42) was obtained for *slop*_*detachmentRate*_ = 4 *s*^−2^ and *intercept*_*detachmentRate*_ =-1.6 *s*^−1^. B) The depolymerization and severing rates were dramatically increased (at least two orders of magnitude) and all filaments were kept. The best fit was obtained for [Cofilin] = 175 *μM* and severing rate = 1 *s*^−1^, depolymerization rate constant = 10 *s*^−1^, 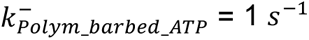 = 1 *s*^−1^ (*w* = 0.14). A and B) Blue: simulations; Red: experimental data for Act1p (adapted from Lacy et al., (2019), OB: Old Barbed end, NB: New Barbed end, OP: Old-Pointed end, and NP: New-Pointed end.

Another possible explanation for the shorter average dwell time is that the rate constants for filament severing or depolymerization are markedly higher *in vivo* than *in vitro*. Indeed, filament tension, compression, twisting, and bending promotes their severing (Huehn et al., 2018; McCall et al., 2019; Murrell & Gardel, 2012; Schramm et al., 2017; Wioland et al., 2019), which is enhanced by crosslinking (Breitsprecher et al., 2011; Ma & Berro, 2019; Mak et al., 2016; McCall et al., 2019). Since yeast CME requires large forces to overcome the high turgor pressure, the actin filaments around the endocytic pits are likely under high mechanical stresses and consequently, their severing rate could be larger than observed *in vitro*. In addition, proteins such as Twinfilin, Aip1, and SRV2/CAP enhance filament severing and/or their depolymerization rates (Balcer et al., 2003; Moseley et al., 2006; Romet-Lemonne & Jégou, 2021; Schramm et al., 2017; Shekhar et al., 2019). To account for these effects in our model, we tested larger ADF/cofilin binding, severing, and depolymerization activities. We varied the parameters to obtain the best fit between simulated and experimental dwell time distributions, but to avoid overfitting we limited the number of optimization iterations (**Fig. 2**B).

Fast actin disassembly notably reduced the average dwell time and produced simulated dwell time distributions for actin subunits very similar to the experimental distributions (**Fig. 2**B). While the *w* between the experimental distribution and the simulated distributions assuming the removal of severed filaments from the actin network was 0.42, simulations with fast actin disassembly led to a much smaller *w* value of 0.14. For this reason, we used the fast actin disassembly model for the rest of our simulations. However, to avoid overfitting, we did not exactly use the set of parameters that minimize the *w* but instead, we randomly picked a set of values with an effect size, *w* = 0.16, close to the minimum value (**Table 2**).

Under the fast actin disassembly assumption, even though each subunit resided in a filament for only a short time (∼1.5 *s*), the filaments’ lifetime was 7 ± 2 *s*, while almost one-third of filaments lasted for the entire simulation (10 *s*, **Fig. S4**). The length of filaments was 49 ± 43 *nm* (Ave ± SD, n=13787, range=6-350 *nm*, **Fig. S2**A and C), consistent with values measured experimentally, ∼50 *nm* (Young et al., 2004), and at the plasma membrane, ∼100 *nm*, (Rodal et al., 2005) or estimated using quantitative microscopy and mathematical modeling, ∼50-150 *nm*, (Berro et al., 2010; Berro & Pollard, 2014b; Sirotkin et al., 2010). In contrast, using the nominal slow disassembly rates led to much longer filaments after 10 seconds (420 ± 80 *nm*, **Fig. S2**B and D). While this analysis considered only the filaments that contained the old barbed ends after each severing event (i.e. the filaments that are the most likely to remain attached to the endocytic structure *in vivo*), we repeated our analysis including all the fragments generated during the simulations. When disassembly was fast, the ensemble average rapidly converged within 6 seconds, whereas simulations using nominal slow disassembly rates converged in ∼100 seconds (**Fig. S2**D). These simulations also allowed us to determine the average number of uncapped fragments generated, as these filaments play a crucial role in force generation during CME. Notably, after a 10-second simulation using fast disassembly rates, each filament generated ∼40 other uncapped filaments on average (**Fig. S2**C). This finding are consistent with the results reported by Wang et al. (2016).

### Release of severed fragments alone is not sufficient to explain the dwell time distributions of capping proteins

Next, we aimed to determine the dwell time distribution of a barbed end capping protein and compare it with the experimental distribution for the Acp1p subunit of the canonical capping protein (the Acp1p/Acp2p heterodimer) reported in Lacy et al., (2019). We first hypothesized that the dwell time of a capping protein is sensitive to the assumptions we make about the fate of both fragments generated after a severing event (hereafter called old-barbed end and old-pointed end fragments). Indeed, while discarding a fragment affects the dwell time distribution of actin subunits, its impact on the capping protein dwell time distribution may be more pronounced because capping proteins are present only at the barbed end of a filament and not homogeneously distributed like tagged actin subunits are (**Fig. S5**). Since capping proteins have a very slow dissociation rate constant from filament barbed ends (∼ 10^-3^ *s*^−1^), during the short course of an endocytic event (∼10 seconds) it almost exclusively dissociates from the filament after the barbed end disappears from the actin-network either due to filament depolymerization or to filament detachment from the actin network. Even though filaments at endocytic sites are heavily interconnected by fimbrin crosslinkers, a fragment severed from an actin filament may be released into the cytoplasm. We tested four different hypotheses: i) always discard the old-barbed end fragment (**Fig. S6**A), ii) discard the old-pointed end fragment (**Fig. S6**B), iii) discard either the old-barbed end or the old-pointed end fragments with equal probability (**Fig. S6**C), iv) discard either the old-barbed end or the old-pointed end fragment with a probability inversely proportional to their length (i.e. following a Bernoulli process with probability p/(1-p), where p is proportional to the inverse of the fragment length) (**Fig. S6**D). We limited ourselves to these four specific hypotheses as we have covered the general detachment of a filament from the actin network in the previous section. Our simulations showed that systematically removing the old-barbed end or old-pointed end fragments (hypotheses i and ii) or removing fragments with a length-dependent probability (hypothesis iv) did not produce good fits. Removing either fragment with equal probability (hypothesis iii) produced a good fit with the dwell time distribution for actin but not for the capping protein. The only hypothesis that produced excellent fits to the experimental dwell time distribution for actin and the capping protein was when both fragments were kept during the simulation (**Fig. 3**A, **Fig. S5** and **Fig. S6**E), which is the assumption we will keep for the rest of this study. This result suggests that the simulated filament can be considered well-connected to the network, which is consistent with experimental data showing actin filaments are heavily interconnected by fimbrin crosslinkers (Berro & Pollard, 2014a; Picco et al., 2018; Sirotkin et al., 2010) and validates the assumption used in previous sections.

**Figure 3.**
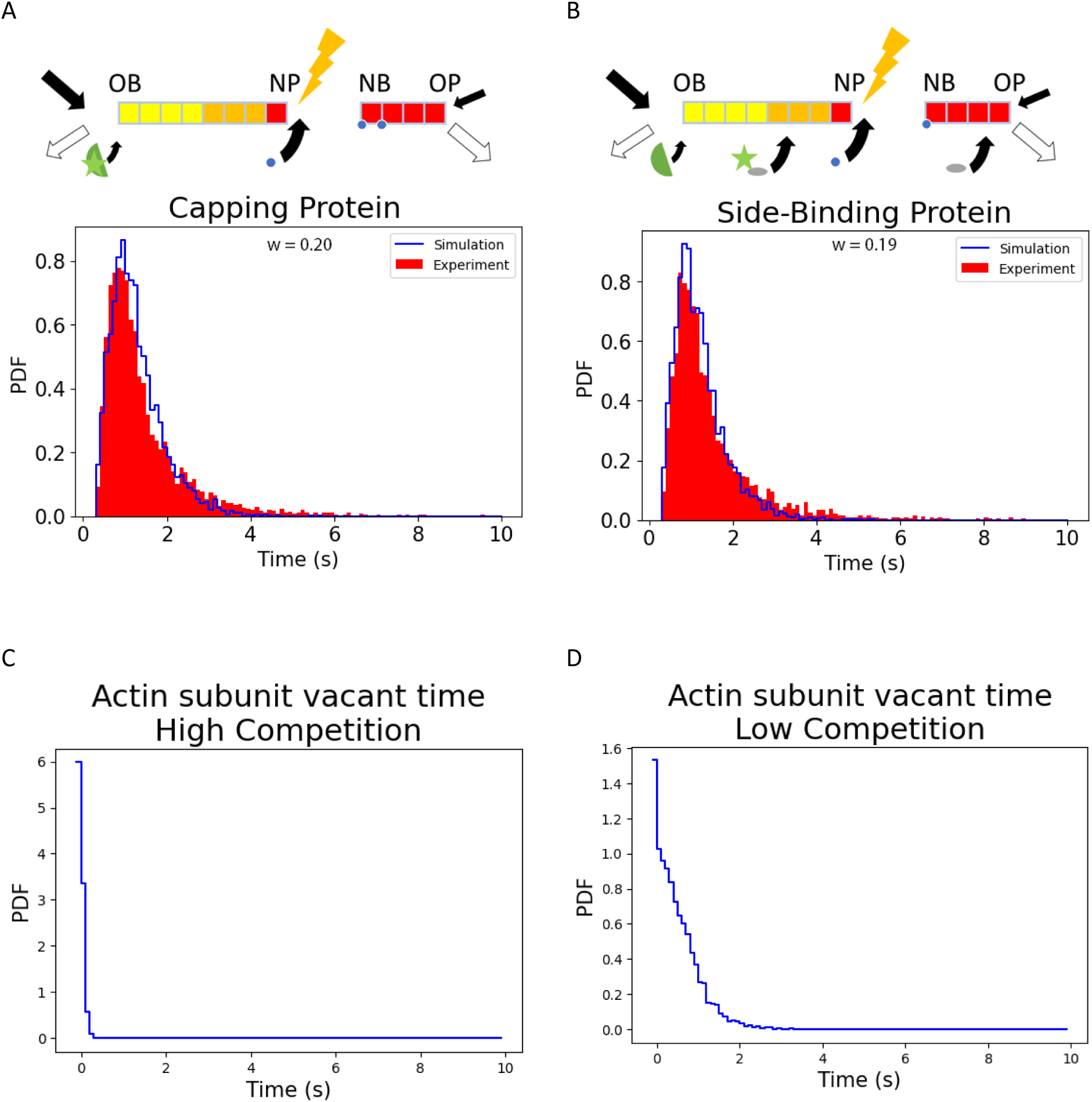
Dwell time distributions of capping and side-binding proteins. A) Distributions for the barbed-end capping protein. We used the optimum parameters as listed in **Table 2**, *w* = 0.20. Blue: simulations (adapted from Lacy et al., 2019); Red: experimental data for the capping protein. B) Distribution for a representative side-binding protein. Fast binding of other proteins to the side of the actin filament is required to saturate the binding sites and obtain distributions that fit the experimental data well, *w* = 0.19. Blue: simulations; Red: experimental data for Myo1p (adapted from Lacy et al., 2019). C and D) Distribution of vacancy times of subunits of actin filaments for high competition, 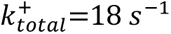 (C) and low competition, 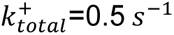 (D) between actin side-binding proteins, where 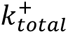 is the summation of association rates of all of the side binding proteins competing over an available binding site.

### Side-binding sites on the filament are occupied quickly

In Lacy et al., (2019) dwell time distributions of proteins that bind to the side of actin filaments such as type-I myosin Myo1p, HIP1R homologue End4p, and the Arp2/3 complex are similar to the distributions for actin subunits. Even though some of these proteins may interact not only with actin subunits but also with the plasma membrane or other proteins, the similarity between actin and side-binding protein dwell time distributions made us first hypothesize that they are all directly determined by actin filament dynamics. However, this hypothesis requires that the association rates of side-binding proteins are much faster than the typical dwell time of actin subunits. If they were not, the actin filament would be sparsely decorated by side-binding proteins and any new protein could bind virtually anywhere along the filament. Their dwell time would therefore be significantly shifted towards smaller values. Therefore, for a side-binding protein to have a dwell time distribution like actin’s dwell time distribution the binding rates of the side-binding proteins must be fast. Consequently, any actin subunit becomes occupied shortly after polymerization, and filaments are heavily decorated with side-binding proteins.

To test this hypothesis, we assumed that each actin subunit possesses a single binding site for side-binding proteins and that this binding site is independent of the ADF/cofilin binding site (Dominguez & Holmes, 2011). In addition, to simplify the model and without loss of generality, we assumed that two different populations of proteins compete over the available sites for side-binding proteins (one population is the protein of interest, and the second population contains all the other proteins competing with the protein of interest over the available binding sites). As a first approximation, we assumed that the dissociation rates of both side-binding proteins and other competing proteins are zero and we screened possible values for their association rates (**Fig. 3**B). Fits between experimental and simulated dwell time distributions improved with increasing association rates. The best fit was obtained when the association rates for the protein of interest and the competitor were 3 s^-1^ and 15 s^-1^ respectively (**Fig. 3**C, **Fig. S7**A. *w* =0.18). In this case, the average vacant time of binding sites on actin subunits was 0.05 ± 0.05 seconds (mean ± std). After we removed the competitive side-binding proteins from the simulation, the average vacant time increased to 0.5 ± 0.5 seconds (mean ± std) and the fit between distributions dramatically declined (**Fig. 3**D & **S7**B, *w* = 0.87). Therefore, our model demonstrates that side-binding proteins decorate actin subunits shortly after they are polymerized. Although it would be difficult to validate this prediction directly through experiments because it would require measuring many *in vivo* protein concentrations and *in vitro* binding rate constants that are currently unknown, we can estimate the actin subunit’s vacant time using typical values from previously reported concentrations and association rate constants of actin-binding proteins. There are ∼20 endocytic proteins that can bind to the side of an actin filament (Goode et al., 2015; Lacy et al., 2018) with cytoplasmic concentrations ∼1-4 *μM*, and association rate constants ∼0.1-10 *μM*^−1^*s*^−1^ (Arasada & Pollard, 2011; Berro et al., 2010; Berro & Pollard, 2014b; Chen & Pollard, 2013; Gonzalez Rodriguez et al., 2023; Picco et al., 2015; Sirotkin et al., 2010; Sun et al., 2019). Using these numbers, we estimate that the vacant time is in the range of 0.001-0.5 s range which agrees well with the results of our simulations.

### The two peaks in fimbrin dwell time distribution are not due to different populations of fimbrin from different cellular processes

Compared to actin and other actin-associated proteins, the dwell time distribution of the actin filament crosslinking protein fimbrin has the particularity of containing two peaks (Lacy et al., 2019)-a peak around 1.5 seconds comparable to the peak present in the distributions of actin and actin-binding proteins (the “slow peak”) and an extra peak around 0.5 seconds unique to the fimbrin dwell time distribution (the “fast peak”). First, we wondered whether these short-lived events could be due to fimbrin molecules associating with non-endocytic structures such as the cytokinetic ring, actin cables, or other actin structures, instead of fimbrin molecules within endocytic patches. Previous reports of fluorescently tagged fimbrin indicated that it primarily localizes to endocytic patches (Christensen et al., 2019; Goode et al., 2015; Picco et al., 2015; Sirotkin et al., 2010; Wu & Pollard, 2005) but a small fraction may also associate to the contractile ring and actin cables (Li et al., 2016; McDonald et al., 2017; Skau et al., 2011; Wu et al., 2001). It is also possible that our single-molecule tracking might have been sensitive enough to detect rare non-CME localization which would have been difficult to observe in cells where fimbrin was GFP-tagged due to the overwhelming brightness of endocytic patches.

To test the hypothesis that the fast peak could be due to fimbrin molecules outside of endocytic patches, we analyzed single-molecule tracks from a new set of movies of Fim1p-SiR labeled cells, recorded as in Lacy et al., (2019), by grouping the tracks based on their location in each cell. We semi-automatically assigned tracks in two groups: the cell tips, defined as the regions covering the 25% of the cell closest to the tips, or in the middle of the cell, defined as the region that covers 50% in the middle of the cells. Endocytic sites are located throughout the cell but are concentrated at the cell tips during interphase (Berro & Pollard, 2014a; J. Marks et al., 1986; J. R. Marks & Hyams, 1985), whereas cytokinetic rings assemble at the midpoint of dividing cells where endocytic sites are also concentrated during cytokinesis. If the short-lived fimbrin events were due to non-endocytic structures, they would be less prominent among tip-localized patches than in the middle of the cells. However, we found no noticeable difference between the dwell time distributions for tracks localized in the middle (*w* = 0.19) or at the tip (*w* = 0.11) of the cell (**Fig. 4**A and B). Both groups retained the fast peak around 0.5 seconds in the same relative proportion, suggesting that these short-lived events are not uniquely attributable to other cellular structures and do indeed occur within endocytic patches. This result corroborates our previous experimental data showing that virtually all single-molecule fimbrin events colocalized with endocytic patches where Acp2p-GFP was also detected (Lacy et al., 2019).

**Figure 4.**
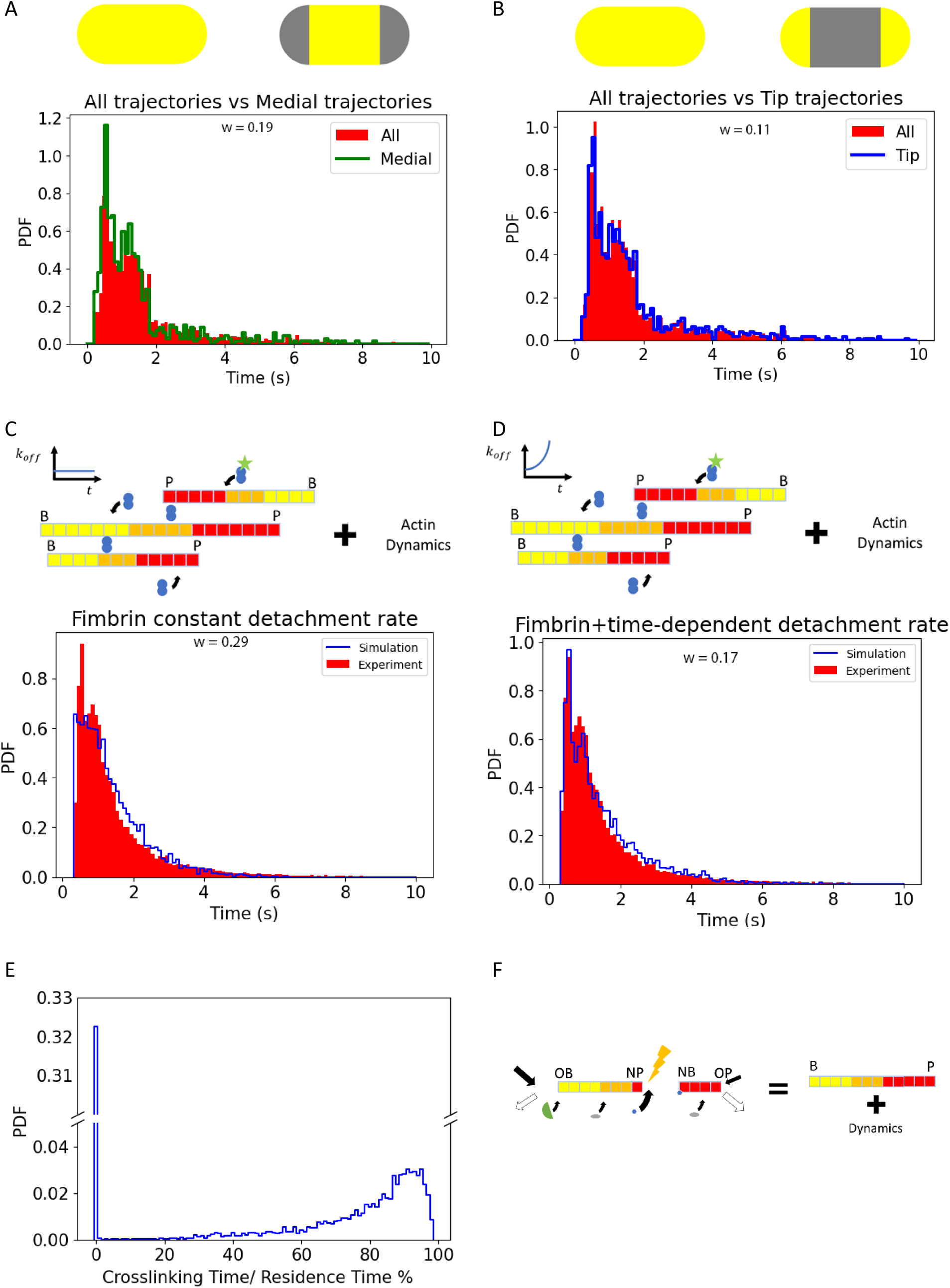
The second “fast” peak in fimbrin dwell time distribution can be explained by a force-dependent detachment of fimbrin. A and B) Experimental data for Fim1p dwell time distributions were pooled depending on their location in the cell: (A) middle part of the cell, *w* = 0.19, (B) tip of the cell, *w* = 0.11. Both the slow and fast peaks are present in the two subsets, suggesting the fast peak (∼0.5 *s*) is not caused by a different process such as cytokinesis, and is ubiquitous in different regions of the cell. C) Using a fixed off-rate constant for the fimbrin cannot explain the existence of the fast peak, *w* = 0.29. D) When the detachment rate of fimbrin linearly increased over time upon crosslinking two filaments, the simulated dwell time distributions contain two peaks, *w* = 0.17. E) Distribution of time-fraction a fimbrin molecule is engaged in crosslinking two filaments. While almost 30% of the fimbrin molecules dissociate before crosslinking two filaments (first bin), the fimbrin molecules that do crosslink filaments remained bound to at least one filament 79±17% of the time on average. F) The compacted schematic includes all dynamics of the actin filaments as described before

### The existence of two peaks in the fimbrin dwell time distribution cannot be explained by simple mass-action kinetics of a protein with two actin-binding sites

Next, we investigated whether the sub-population of short-lived fimbrin tracks could be explained by fimbrin’s complex binding kinetics. Because each fimbrin molecule can bind two actin filaments with its two actin-binding domains (ABDs), it can be in three different states: i) diffusive when a fimbrin molecule is not bound to any filament, ii) one-side bounded, when only one ABD is associated to a filament, iii) crosslinking, when both ABD1 and ABD2 are associated to two filaments and crosslink them. Because of these three states, the dwell time of the fimbrin is determined by a combination of the attachment and detachment of each ABD from the filaments and the underlying kinetics of actin subunits bound by each ABD. Different scenarios can lead to the apparent dissociation of a fimbrin from the endocytic site. For example, a) a fimbrin molecule is bound to one filament and dissociates before its second ABD binds a nearby filament, b) both fimbrin ABDs of a fimbrin molecule crosslinking two filaments dissociate from their respective filaments, c) the actin subunits to which each fimbrin ABD are bound are disassembled from the filament meshwork, etc.

Since fimbrin molecules crosslink two actin filaments, we ran simulations with two independent filaments (each modeled with full dynamics of polymerization, depolymerization, and severing as described above). For simplicity, we did not explicitly account for any geometrical constraints and the relative position of filaments but implicitly included this information by using different rate constants for the first and the second ABD that binds a filament. In an actual actin network, once one of the fimbrin’s ABD associates with one filament, the other ABD’s motion is constrained, and as a result, it can only bind to an actin subunit on another filament in its proximity. Therefore, we expect to have two different apparent association rate constants based on the state of the fimbrin (diffusive or one-ABD bound). To investigate this hypothesis, we used the two different values for association rate constants of each ABD (the diffusive and one-side-bounded) and we used the same dissociation rate constant for both ABDs independently of the binding state of the fimbrin. The dwell time of one fimbrin molecule was calculated as the time between the attachment of its first ABD and the time of the complete detachment of the fimbrin molecule by unbinding or because the filaments disassembled. If only one ABD dissociates or the actin monomer is bound to be disassembled from the filament, this ABD can associate again with a new actin subunit, as long as the other fimbrin’s ABD remains attached to its filament. Consequently, each ABD might associate and dissociate several times before complete dissociation of the fimbrin from the actin meshwork. We screened a range of values for the fimbrin’s association and dissociation rate constants over one order of magnitude but were unable to find any set of parameters that generates a distribution with two peaks similar to the experimental distribution (**Fig. 4**C). This result suggests our model is incomplete and fimbrin’s kinetics at endocytic patches is more complex than mass action.

### Mechanosensitive kinetics can produce two peaks in the dwell time distributions

When both ABDs of a fimbrin molecule are attached to two filaments in an actin meshwork, the tension and torsion on the fimbrin molecule increase over time (Ma & Berro, 2018), which would accelerate the dissociation kinetics of the fimbrin molecule if one assumes that ABD binding to actin filaments is slip-bond. Based on the result of Ma & Berro (2018) the corresponding potential elastic energy stored in fimbrin has a distribution reminiscent of a Gamma distribution. Our simulations do not explicitly account for the effects of mechanics so we modified the detachment rate of fimbrin ABDs with a phenomenological law such that upon crosslinking two filaments the ABDs detachment rates gradually increase over time according to Bell’s Law 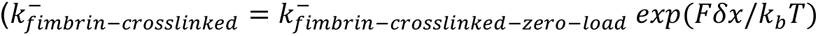, where *F* is the applied force, *δ*x** is the displacement, 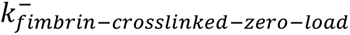 is the dissociation rate constant with zero load, *k*_*b*_ is Boltzmann’s constant, and *T* is the temperature). As a first approximation, we assumed that the potential energy of an ABD increases linearly with time. After varying these parameters over several orders of magnitude, we found that a rapid increase in the dissociation rate leads to the appearance of the “fast” peak in the fimbrin’s dwell time distribution (**Fig. 4**D, **Table 2**). Taken together, our simulation results indicate that the presence of two actin-binding domains on fimbrin, actin filament dynamics, and a force-dependent ABD dissociation rate behavior can recapitulate the experimentally observed fimbrin dwell time distribution at endocytic patches.

Our simulations also allowed us to determine the fraction of time a fimbrin molecule crosslinks two filaments (**Fig. 4**E). Our result indicates that 30% of the fimbrin molecules dissociate without crosslinking filaments. This is caused by high competition over available binding sites as well as highly dynamic actin filaments. Those fimbrin molecules that do crosslink filaments remain attached to both filaments for 79±17% of their time within the actin network.

### Validation of the model’s assumptions and predictions

The minimum number of required ADF/cofilin-bound actins to sever a filament are still under debate (Andrianantoandro & Pollard, 2006; Bibeau et al., 2021; Bobkov et al., 2006; De La Cruz & Gardel, 2015; Elam et al., 2017; Hocky et al., 2021; McCullough et al., 2008; Tanaka et al., 2018). The model we present in this manuscript considers that severing requires at least two consecutive ADF/cofilin bound to the actin filament. To support this hypothesis, we also considered the case where a single ADF/cofilin is sufficient to sever the filament. Under this assumption, while the dwell time distribution of the actin subunits remained almost unchanged (*w* = 0.16, **Fig. S8**A), the distribution of the capping proteins drastically changed (*w*=0.41, **Fig. S8**B). Therefore, our simulations and experimental data support the hypothesis that a single ADF/cofilin is not sufficient for filament severing.

Our model predicts that severing and fast disassembly are critical for short dwell time distributions. Removing or significantly reducing ADF/cofilin from yeast cells is lethal. Therefore we recorded and analyzed movies of the ADF/cofilin mutant allele cof1-M2, whose *in vitro* severing rate in absence of any other factor is up to ∼3 times smaller than wild-type cofilin (Chen & Pollard, 2011). The dwell time distribution of capping protein in the cof1-M2 background was similar to wild-type (**Fig. S9**A). However, simulations using a severing rate three times smaller than our standard conditions showed a significant difference (the average dwell time increased to 1.7 ± 1.1 *s* and *w* = 0.5, **Fig. S9**B). These results support the predictions of our model and previous experimental data showing the severing rates are largely enhanced by mechanical stress and synergies between ADF/cofilin and other proteins such as Twinfilin, Aip1, and SRV2/CAP (Huehn et al., 2018; McCall et al., 2019; Murrell & Gardel, 2012; Schramm et al., 2017; Wioland et al., 2019).

Our simulations did not include the Arp2/3 complex. While the Arp2/3 complex is critical to nucleate new actin filaments, it marginally affects filament disassembly, since most pointed ends are created by filament severing and are not capped by an Arp2/3 complex. To verify this rationale, we performed simulations where all the filaments were nucleated with an Arp2/3 complex at their pointed end. With the rate constants of our best model, the dwell time distributions remained qualitatively the same, and differences could be compensated by increasing the depolymerization rate of the barbed end, and by increasing the severing rate by a factor of two (**Fig. S10**, *w* =0.17). Therefore, the Arp2/3 complex has a minimum effect on the overall results of our simulations.

## Discussion

Previous conceptual models of CME have typically assumed that the membrane coat and actin meshwork assemble throughout the endocytic event and then disassemble after the vesicle is released. However, our group recently showed that actin and other endocytic proteins are rapidly turned over during an endocytic event in wild-type cells with dwell times of around 1 to 2 seconds (Lacy et al., 2019), confirming predictions from previous mathematical modeling (Berro et al., 2010). These data demonstrated that the endocytic machinery is much more dynamic than was previously appreciated. It is also reminiscent of other sub-cellular processes based on actin, which are under high turnover (e.g., lamellipodia of crawling cells or growth cones, assembly and constriction of the cytokinetic ring, focal adhesions). Using stochastic simulations of the dynamics of actin filaments and actin-binding proteins, we demonstrate here how the unique kinetics of actin and actin-binding proteins can generate dwell time distributions similar to the ones obtained experimentally.

### Predicted kinetics for actin and actin-binding proteins

Our simulations showed that the peaked shape of actin monomer dwell time distributions can be achieved through a combination of filament polymerization, filament severing (e.g., by ADF/cofilin), and fast-pointed end depolymerization. This result is compatible with recent *in vitro* evidence of the synergistic effect of ADF/cofilin, SRV2/CAP, and other proteins like Aip1 to disassemble filaments at rates in the order of 90 subunits/s per pointed end (Shekhar et al., 2019). Previous mathematical modeling predicted that fast disassembly of actin filaments is required to explain the experimentally measured evolution of the number of actin molecules at endocytic sites (Berro et al., 2010). In that study, the release of severed filaments was favored over very fast pointed-end depolymerization because the required depolymerization rate required to fit the experimental data (83 subunits/s for each pointed end) was orders of magnitude larger than any rate constant reported at the time. This fast depolymerization rate constant has since been corroborated by recent experimental *in vitro* data (Shekhar et al., 2019; Wioland et al., 2017) and even produces better fits to our single molecule dwell time distribution of actin and actin-binding proteins in our new model presented in this paper. Our model predicts that mutations on proteins such as Srv2/CAP that reduce their actin depolymerization activity or reduction of the tension on the filaments should have a large effect on the actin dwell time distribution, whereas ADF/cofilin mutants with lower severing activity should have a much milder effect on those distributions.

Our model also predicts that actin subunits are decorated by actin-binding proteins shortly after polymerization (∼0.05 *s*). Because the detachment half-time of known actin side binding proteins is significantly slower (∼10-1000 *s*) than the lifetime of actin filaments (∼ 1 *s*), a consequence of this prediction is that the filament side is fully decorated by actin-binding proteins. Since endocytic sites contain numerous proteins that have actin-binding domains, full filament decoration would be possible even if their cytoplasmic concentrations and binding rate constants are moderate. We also predict that protein affinity has less impact than its binding rate (i.e., the product of the protein concentration and its binding rate constant) on its relative occupancy of endocytic actin filaments compared to other actin-binding proteins - the larger the protein binding rate, the larger its occupancy. We also predict that proteins with a slow binding rate will be excluded from endocytic patches. Future quantitative microscopy experiments will aim to determine the occupancy of actin subunits in the endocytic patches.

### Evidence for a force-dependent unbinding of fimbrin

Our new analysis of fimbrin’s dwell time distribution demonstrates that the two peaks, which are unique among the measured endocytic proteins, are not attributable to non-endocytic events or to the fact that fimbrin has two actin-binding domains. Instead, we show that the two peaks are consistent with an increase over time in the tension on fimbrin molecules that crosslink two filaments. Note also that two-color experiment with Fim1-SNAP/Acp2-GFP found over 95% of Fim1-SNAP events colocalized with Acp2-GFP patches (Lacy et al., 2019). Currently, we do not know whether the binding of fimbrin ABDs to actin behaves as a slip-bond or a catch-bond, which has been proposed for other crosslinkers and actin-binding proteins (Huang et al., 2017; Owen et al., 2022; Schiffhauer et al., 2016; A. Wang et al., 2022). Based on the result of (Ma & Berro, 2019) our model favors the slip-bond hypothesis. However, more experimental work will be necessary to test this hypothesis.

### Implications for the mechanics of endocytosis

Our data further demonstrate that the endocytic machinery is highly dynamic and essentially driven by fast actin assembly and disassembly. This property has several impacts on how actin produces force to shape the plasma membrane into a vesicle. Fast turnover allows the actin meshwork to reorganize continuously, allowing it to reorient the force applied to the plasma membrane at different stages of the endocytic process. Fast turnover also allows the actin meshwork to produce forces through a higher-order organization (e.g., by releasing stresses) as was suggested in motility reconstitutions *in vitro* (Dayel et al., 2009). Fast turnover also keeps the system far from equilibrium, which allows the sustained accumulation of elastic energy in filament crosslinkers and its subsequent release into work (Ma & Berro, 2018), or the sustained production of compressive forces by crosslinking (Ma & Berro, 2019), for example.

Our experimental data and mathematical model are also compatible with the idea that proteins that bind both lipids and actin filaments directly via other proteins (e.g., End4p, Myo1p, clathrin) can be pulled by the actin meshwork and stripped from the plasma membrane if they bind more strongly to actin than to lipids. In addition, the pulling of these proteins may create lipid flows which could further help in the formation, elongation, and scission of the endocytic pit.

## Materials and Methods

### Measure of differences between dwell time distributions

There are several measures to evaluate the differences between the two distributions. We chose Cohen’s *w* value as the experimental data were collected on a discrete interval, and features of the distributions such as the existence of the fast peak in fimbrin were important. *w* is defined as the square root of the standardized chi-square statistic, 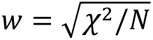. Like chi-square, *w* is sensitive to noise for the bins with small counts. For this reason, we measured the *w* value only for the dwell times from 0.3 to 3 seconds. This interval contains almost 90% of the data points for all the datasets we used in this study.

### Actin filament simulations

We used the rejection sampling Monte Carlo simulation method (Vestergaard & Génois, 2015), an improved version of the *τ*-leaping simulation method, to simulate all reactions. The simulation timestep *τ*, is calculated as 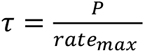, where *P* is a constant (*P* = 0.1), and *rate*_*max*_ is the maximum value of all the rates used in the simulation. As the maximum rates do not change during the simulation, we do not need to update *τ* at each time step.

For each reaction, the model generates a random number, *R*, with a uniform distribution between 0 and 1. If the generated number satisfies *R* ≤ (*rate of the reaction*)*τ*, the model performs the reaction. For the association or dissociation of a subunit, once the reaction is triggered after satisfying the above condition, a new subunit is added or removed, respectively. For every polymerized subunit, there are several possible reactions based on their status as listed in **Table 1**. To decorate a subunit with ADF/cofilin, the subunit must be in ADP state. Cooperative ADF/cofilin binding is implemented by checking the status of adjacent subunits. All the changes applied to the filament and its subunits at the end of each time step as indicated in the diagram in **Fig. 5**.

**Figure 5.**
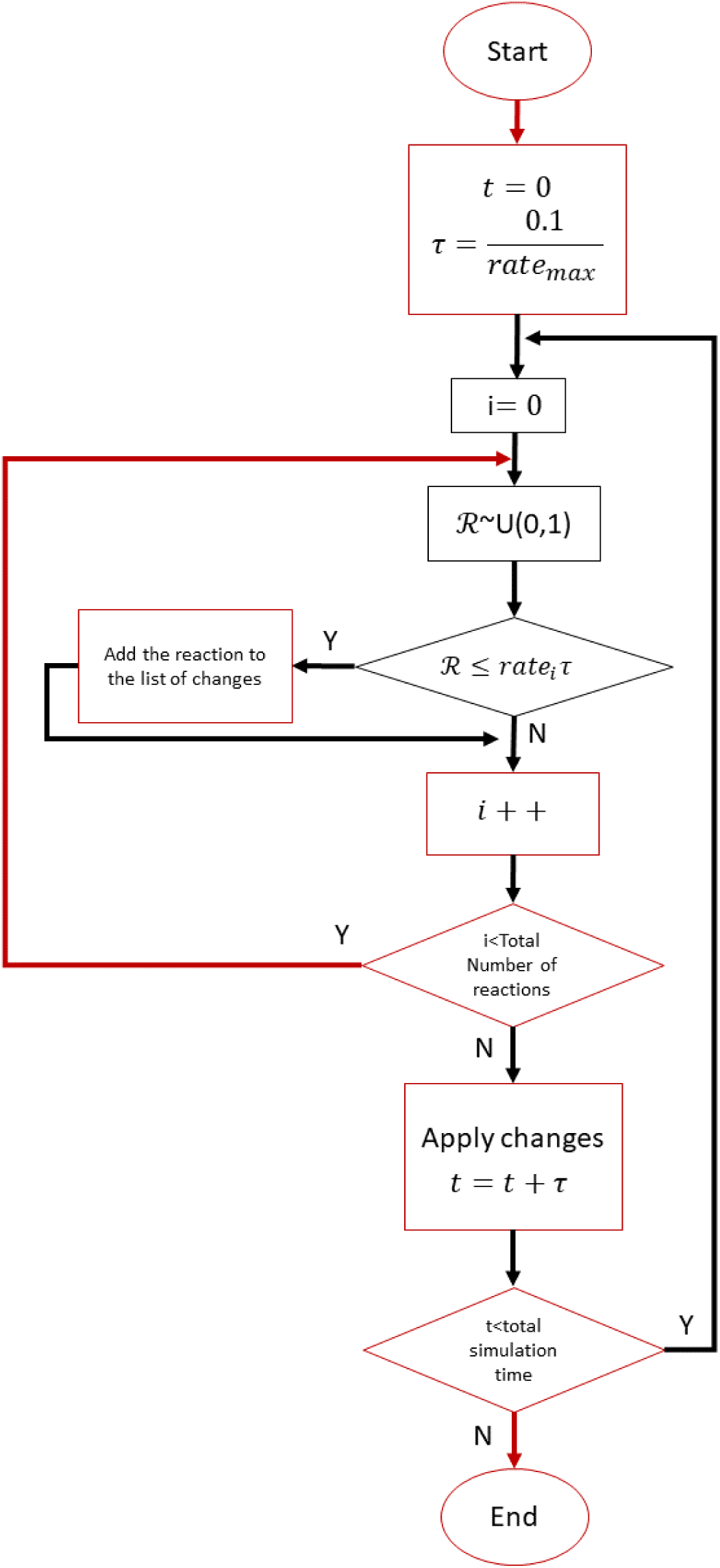
Flowchart of the simulations. The list of reactions consists of the reactions listed in **Table 1** for both the filament(s) as well as all of the subunits. *R* is a random number with uniform distribution between 0 to 1. *τ* is constant and chosen in such a way that *rate*_*i*_ *τ* is smaller than 0.1 for all the reactions. This condition is necessary to obtain accurate results. At the end of each time step, we apply the changes corresponding to the triggered reactions.

For fast actin disassembly, we assumed that actin subunits depolymerize from both ends with higher rate constants (Shekhar et al., 2019; Wioland et al., 2017, 2019). We also assumed that ATP is hydrolyzed at a fast rate (Berro et al., 2010; Ti & Pollard, 2011). Before the onset of the simulation (t=0 second), a four-subunit seed filament is first assembled using the same rate constants used in the rest of the simulation. This was achieved by allowing precisely four subunits to polymerize before time was set to 0. During the simulations, any filament that contained fewer than two subunits was removed. If at any point during the simulation no filament remained, we reinitiated one new seed filament containing four subunits (as describe above) and continued the simulation. After severing of a filament, each one of the two fragments was considered a separate filament.

### Filament removal

It is possible that a filament within the endocytic actin meshwork detaches from the network and diffuses away. To incorporate this feature into our model, we considered the detachment of the filament from the network to be a random process with a reaction rate that might or might not depend on the age of the filament. As a first approximation, we used the following equation to calculate the filament detachment rate *filamentDetachmentRate* = *max*{0, *slop*_*detachmentRate*_ ∗ (*filamentAge*) + *intercept*_*detachmentRate*_}, where *filamentAge* is defined as *filamentAge* = *current time* − *mean*(*subunit polymerization time*). As long as *filamentDetachmentRate* is equal to zero, we do not discard the filament.

### Fimbrin detachment

When only one of fimbrin’s ABD was attached, we assumed its detachment rate constant was a fixed value. When both ABDs were attached (the fimbrin is crosslinking), we assumed that the bonds between the fimbrin ABDs and actin filaments were slip-bond (Ma & Berro, 2019), and ABD1 and ABD2 detachment rate followed Bell’s Law, 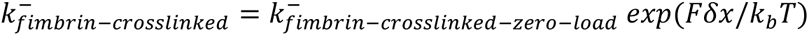, where 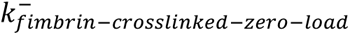 is the dissociation rate with zero-load, *F* is the applied force, *δ*x** is the displacement, *k*_*b*_ is Boltzmann’s constant, and *T* is the temperature. As the force distribution on a cross-linked fimbrin is not known, we assumed the potential energy on a bond followed a Gamma distribution, by previous simulations (Ma & Berro, 2019). We assumed that immediately after the attachment of both fimbrin’s ABDs the tension is minimum and increases over time. To include this in our model, we increase the scale and shape parameter of the distribution linearly. The Gamma distribution was defined as *U* ∼ *Gamma*(*k*, *θ*), where *k* is the shape parameter and *θ* is the scale parameter, *k* = *Shape*_*Slope*_ *t* + *Shape*_*intercept*_, and *θ* = *Scale*_*Slope*_ *t* + *Scale*_*intercept*_.Together these two parameters, *k* and *θ*, determine the average and standard deviation of the distribution.

### Parameter screening

To determine how different parameters affect the model or to find the values that minimize the difference between experimental data and simulation, we used a 2D parameter screening method. A pair of parameters was selected and assigned to the x- and y-axes. For example, to reduce the dwell time of actin subunits, we chose a pair from all actin dynamics-related parameters, e.g., depolymerization rate constants, ATP hydrolysis rates, ADF/cofilin concentration, etc; In **Fig. S3** we have assigned *slop*_*detachmentRate*_ to the x-axis and *intercept*_*detachmentRate*_ for the y-axis. The effect size was calculated for every possible combination of x and y values within the determined discrete range, i.e., For **Fig. S3** A, B, and C, we picked about 100 pairs of values for (*slop*_*detachmentRate*_, *intercept*_*detachmentRate*_). Finally, we ran the simulated for each pair of values and collected the dwell times. The result of each simulation was used to determine *w* value of each tile in the 2D plots. We kept varying the range of the screened parameters till one of the following conditions was satisfied: i) a local minimum value of *w* value emerged within the screening range, ii) further changing the range was not physiologically relevant iii) varying the range of parameters led to small changes of *w* values (less than 0.01).

### Procedure for selecting the optimal values for parameters

To find values that minimize Cohen’s *w*, we used experimentally measured concentrations and rate constants as initial values. Next, we picked pairs of parameters, e.g. (Concentration of ADF/cofilin, severing rate), (random filament removal slope, random filament removal intercept), etc., and screened a range of values for the selected pair as explained previously. We picked the pair of values that minimizes *w* and update the selected parameters. Using the updated values in the last iteration, we picked a different pair of parameters for the next iteration and repeated the screening step as described. It is important to note that by fine-tuning the parameters, it is possible to fit the noise from the experimental data. For this reason, we assumed that the range of all the parameter values that led to *w* smaller than 0.2 were equally valid and stopped the optimization once a *w* smaller than 0.2 was achieved.

### Cell growth, labeling and imaging

Fission yeast expressing Fim1p-SNAP were labeled with SNAP-SiR fluorophore, imaged by partial-TIRF, and analyzed for single-molecule tracking as described in Lacy et al., (2019). Yeast cells were grown in YE5S medium at 32 C till the liquid culture reached the exponential phase (OD of 0.4 to 0.6). After diluting by EMM5S to 0.1 OD, the liquid culture was grown overnight at 25 C. After 12 to 24 h, the culture was diluted to 0.1 OD before adding 1 μM silicon-rhodamine benzylguanine derivative SNAP-SiR647 (SNAP-Cell 647-SiR, New England Biolabs). Culture tubes were wrapped in aluminum foil and incubated on a rotator for 15 hours. Cells were washed 5 times and incubated for 1 more hour on the rotator. Before imaging the cells, the culture was washed 5 more times. It is estimated that 0.1% to 1% of the SNAP-tagged proteins are labeled using this protocol (Lacy et al., 2019).

### Partitioning of the dataset based on the location of the trajectories

To investigate the spatial distribution of events comprising the fast peak of fimbrin dwell times, we partitioned the tracking dataset into subsets based on the location of each trajectory. We manually outlined each cell in the movie. Then, for each trajectory, we determined the location of the track centroid and calculated its distance to the tip of the cell it belonged to. Based on this distance we assigned each trajectory either to the tip-subset or the medial-subset using an adjustable threshold.

## Supplementary Figures

**Figure S1.**
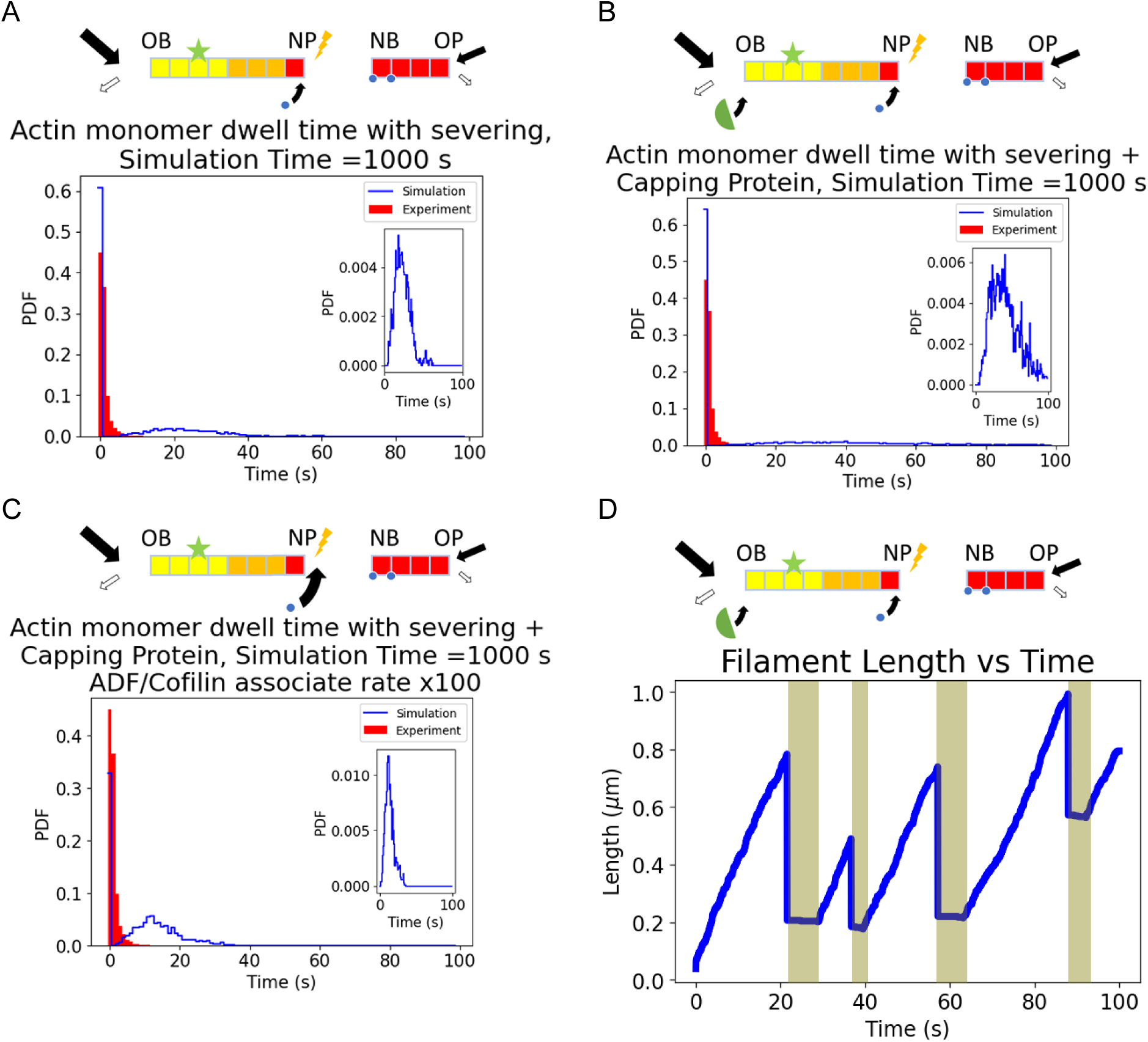
Dwell time distributions of actin subunits with different conditions and simulation time. A) Adding ADF/cofilin introduces a peak at ∼20 s for a long simulation time (1000 s). This peak is absent with a shorter simulation time (Fig. 1B). B) After the introduction of capping proteins, the peak in the actin dwell time distribution is ∼ 40 seconds. C) Increasing the cofilin concentration, its association rate constants, and the severing rate by a factor of ten reduced the peak to ∼10 s. D) Temporal evolution of the length of a representative filament corresponding to **Fig S1**.B. The filament length increased linearly for the first twenty seconds. Then, the length started to fluctuate due to severing. During intervals in gray, the filament shrank as a cluster of cofilactin at the pointed end destabilized it, while the barbed end was capped. After the dissociation of all the cofilactin subunits, the filament elongated again. Insets in A, B, and C: distribution of dwell times omitting the data for the first 10 s.

**Figure S2.**
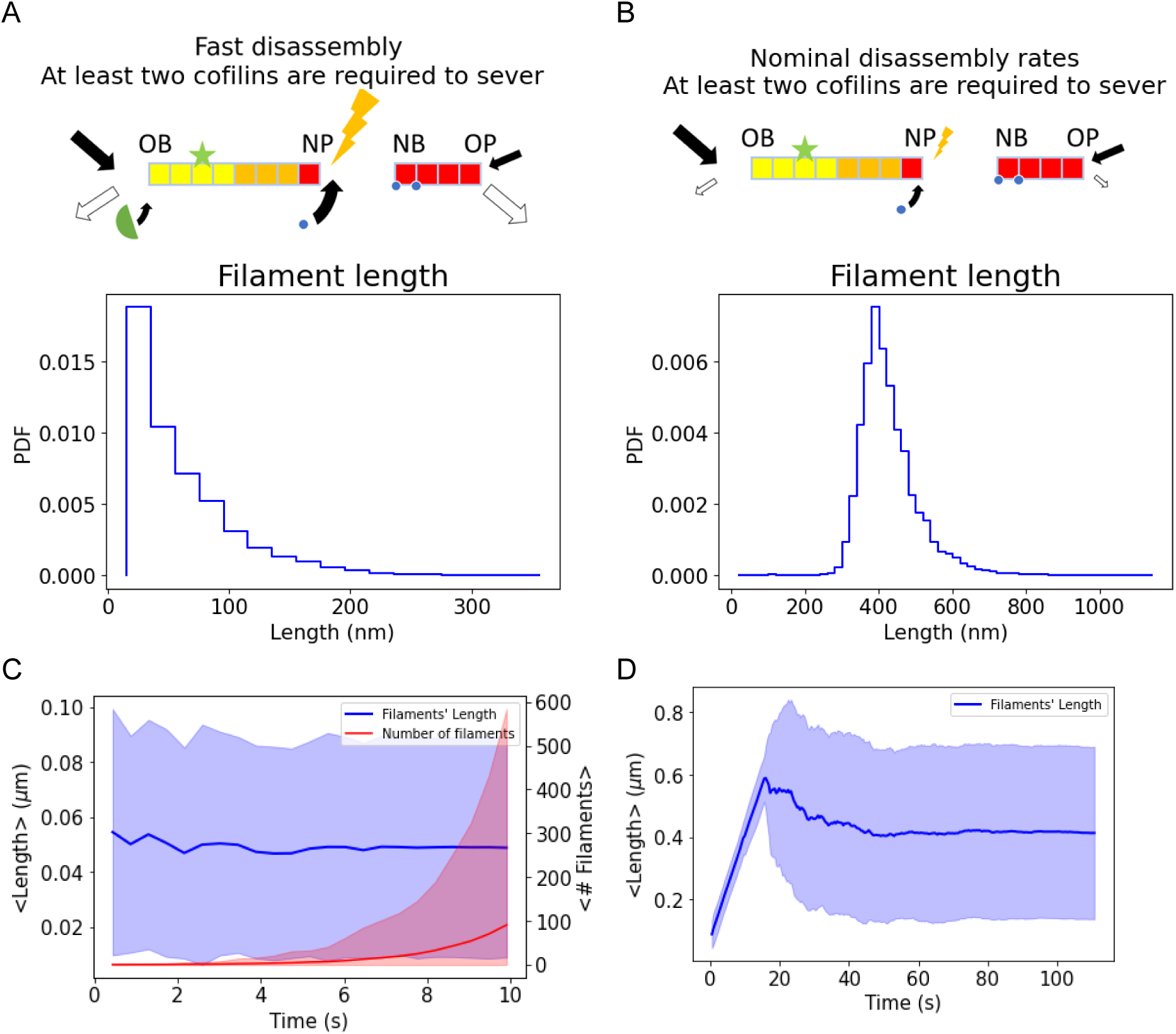
Fast disassembly is necessary to obtain filaments with an average length comparable to the observed actin filaments at CME sites. A and B) Filament length distribution following a 10-second simulation using fast disassembly rates (A), leading to short filaments (49 ± 43 nm), or slow disassembly rates measured *in vitro* (B), leading to long filaments (420 ± 80 nm). Note that in the latter condition (B), the probability of occurrence of short filaments is virtually zero in the first 10 seconds. C and D) Filament length (blue) and number of filaments over time (red, for C only) using fast disassembly rates (C) or slow disassembly rates measured *in vitro* (D). The lines represent averages and the shaded areas represent standard deviations for the filament lengths (blue) in C (73 iterations) and D (43 iterations) or the 95% confidence interval for the number of filaments (red) in C.

**Figure S3.**
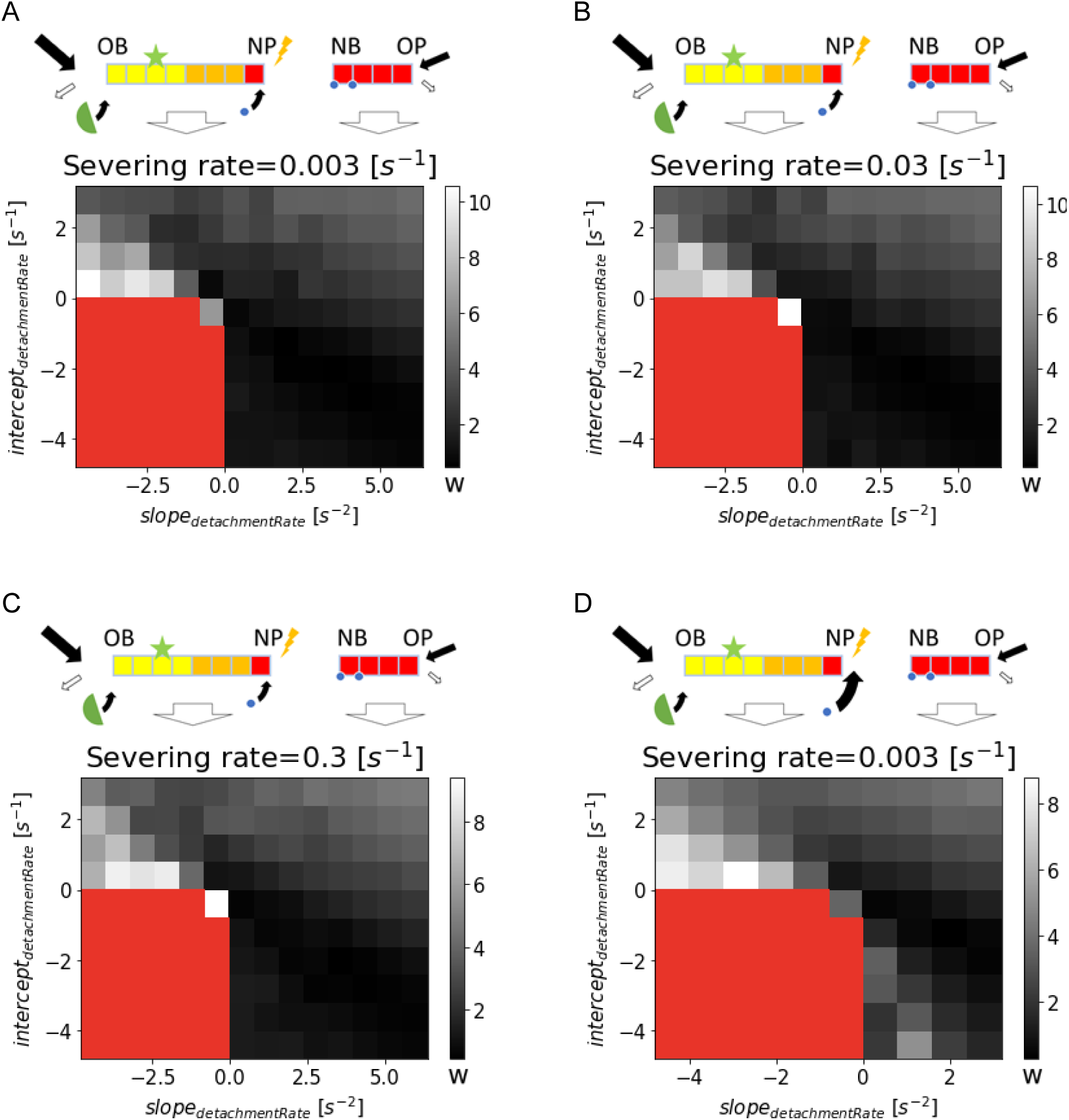
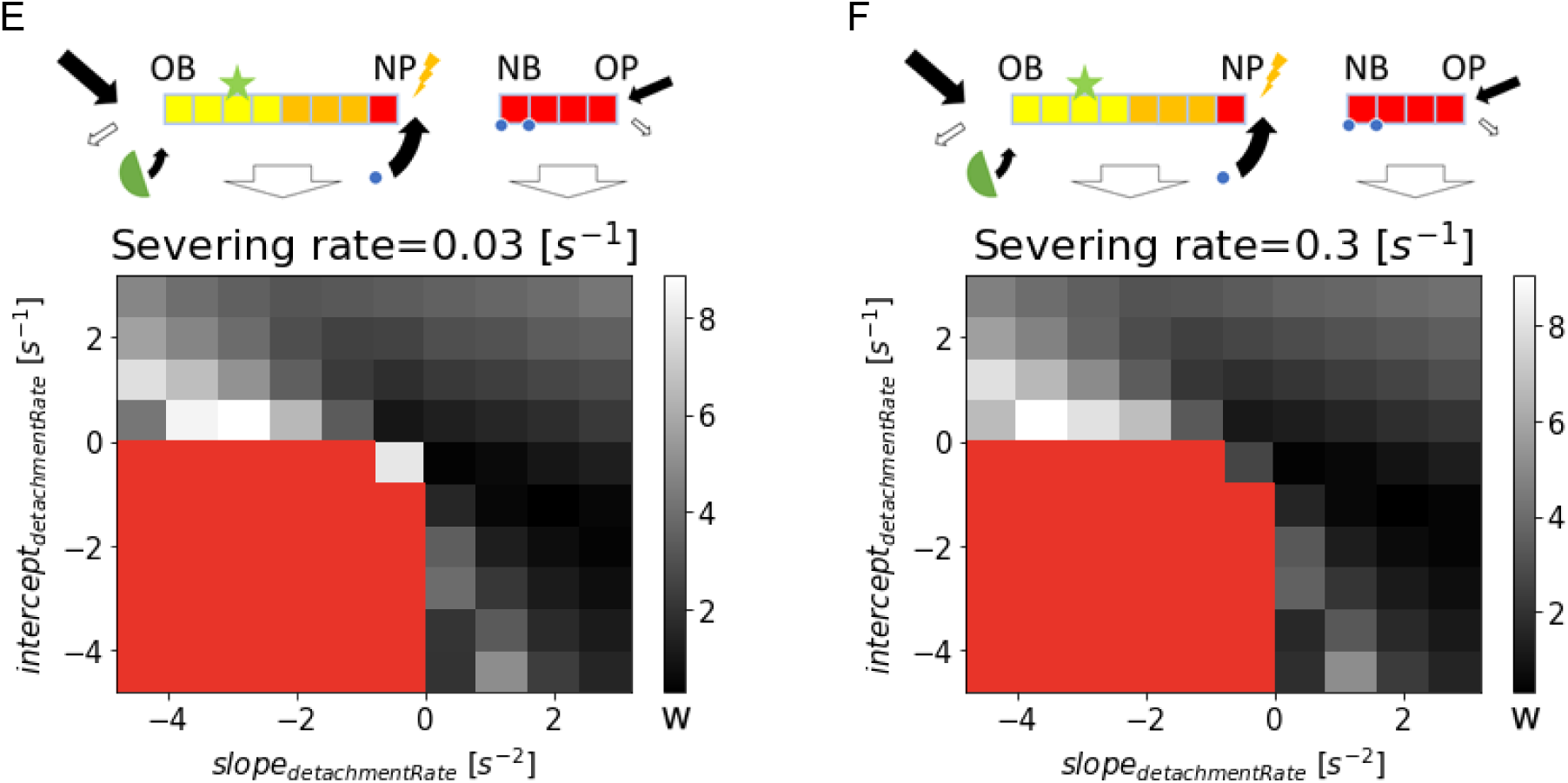
The release of the severed actin filament cannot be responsible for the short dwell time observed in the experimental data – Effect of the concentration of ADF/ cofilin, severing rate constant, and filament detachment rate on the quality of the fit with the experimental data. The heatmap colors correspond to Cohen’s *w*, which measures the difference between the experimental and simulated actin subunit dwell time distributions. Darker gray values correspond to better fits with the experimental data (i.e., lower *w*). Red values correspond to cases where the filament detachment rate is always zero and for this reason, we did not simulate them. We used the reference parameters in **Table 2** except for the concentration of ATP-G-actin for which we used 10 μM to compensate for the effect of applied force on the polymerization rate. The severing rate for A and D is 0.003 s^-1^, for B and E is 0.03 s^-1^, and for C and F is 0.3 s^-1^. The concentration of the ADF/cofilin for A, B, and C is 40 which is equal to the reported concentration of ADF/Cofilin in the cytoplasm (Sirotkin et al., 2010). To evaluate the effect of a higher local concentration of ADF/cofilin for E, F, and G is 150 μM which is equal to the optimum value reported in **Table 2**. The values of *slop*_*detachmentRate*_, *intercept*_*detachmentRate*_ which minimize the *w* value were 4 *s*^−2^ and −1.6 *s*^−1^ for A,B, and C with *w* equal to are 0.44, 0.44, 0.43 respectively, and 2.8 *s*^−2^ and 0.8 *s*^−1^ for E,F and G with *w* equal to 0.29, 0.30, and 0.32 respectively.

**Figure S4.**
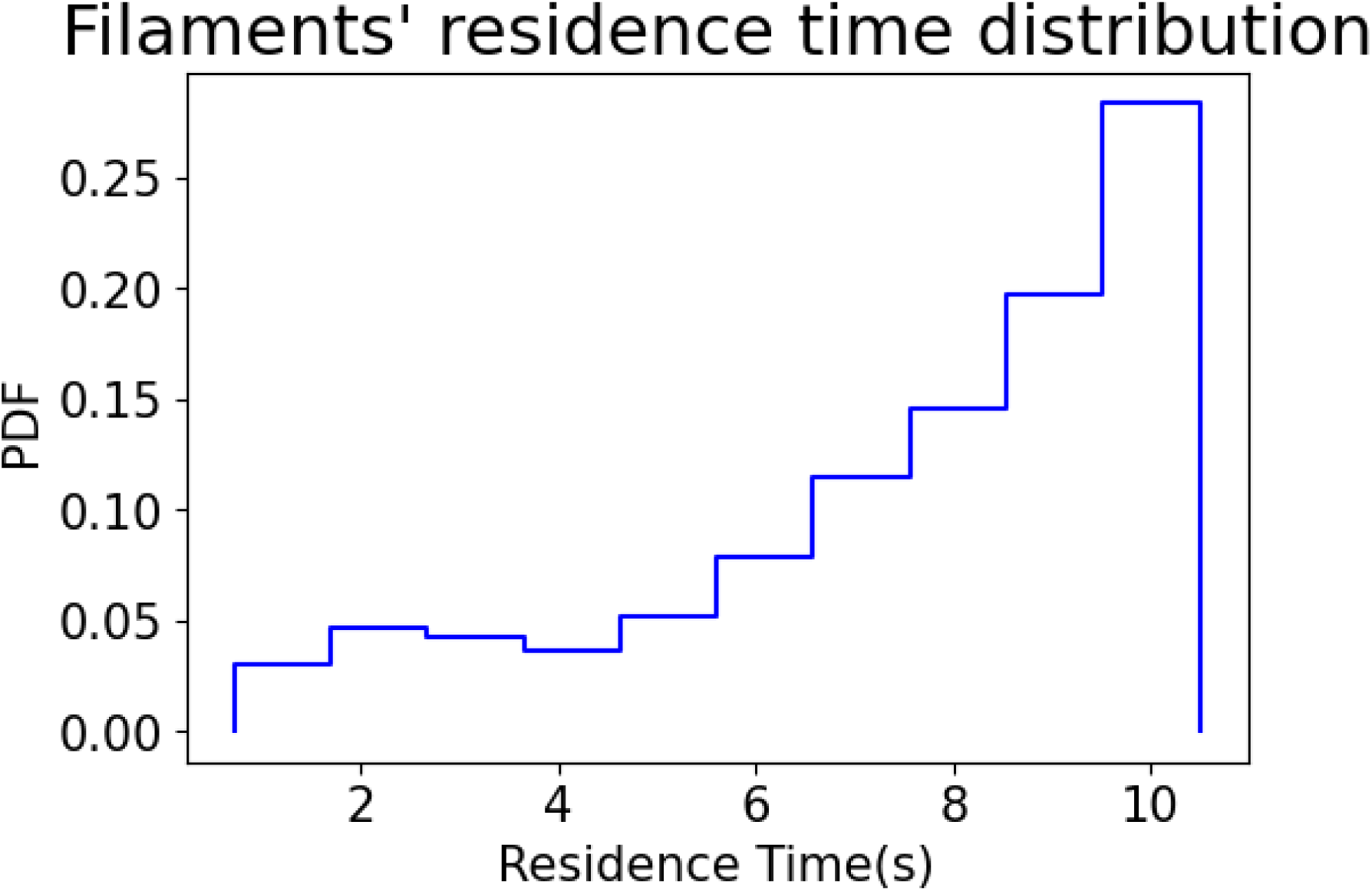
Lifetime distribution of individual actin filaments. While each actin subunit resides inside an actin filament for ∼1.5 s on average (Lacy et al., 2019), almost one third of the filaments do not completely disassemble before the end of the simulation (10 s). A filament lifetime is calculated as the time interval between its creation and either the end of the simulation or its disappearance because it became shorter than 2 subunits.

**Figure S5.**
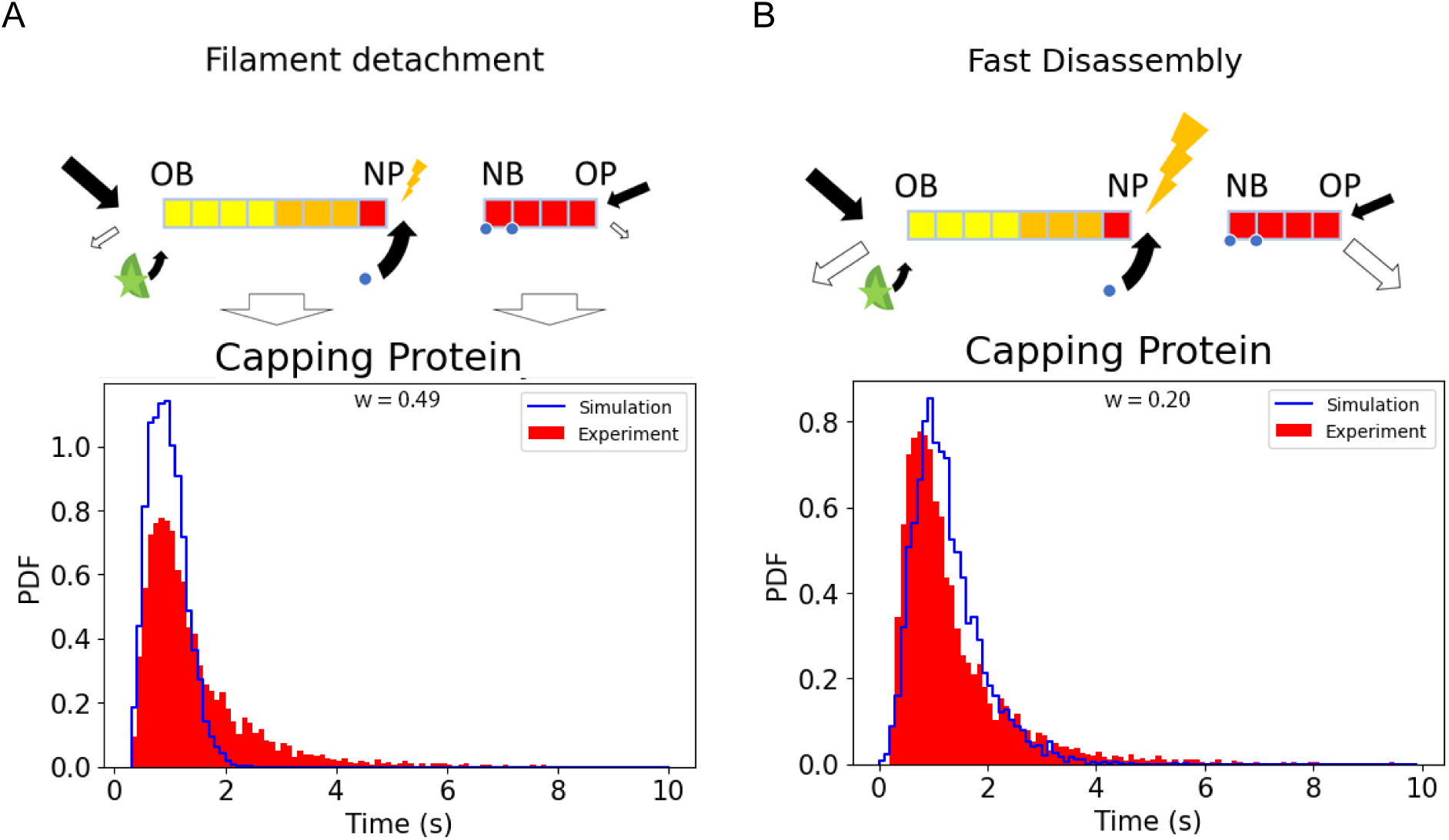
Filament detachment from the actin network leads to pronounced differences in capping protein dwell time. A) Dwell time distribution of capping proteins when filaments can detach and leave the actin network in the same conditions as in Fig. 2A. Although the simulated dwell time distribution of actin subunits is comparable to experimental distributions, the distributions for the capping protein are different (*w* = 0.49). B) Dwell time distribution of capping proteins when filaments are connected to the network and the filament disassembly rate is high, the experimental and simulated dwell time distributions were similar for both actin subunits (Fig. 2B) and capping proteins (*w* = 0.20).

**Figure S6.**
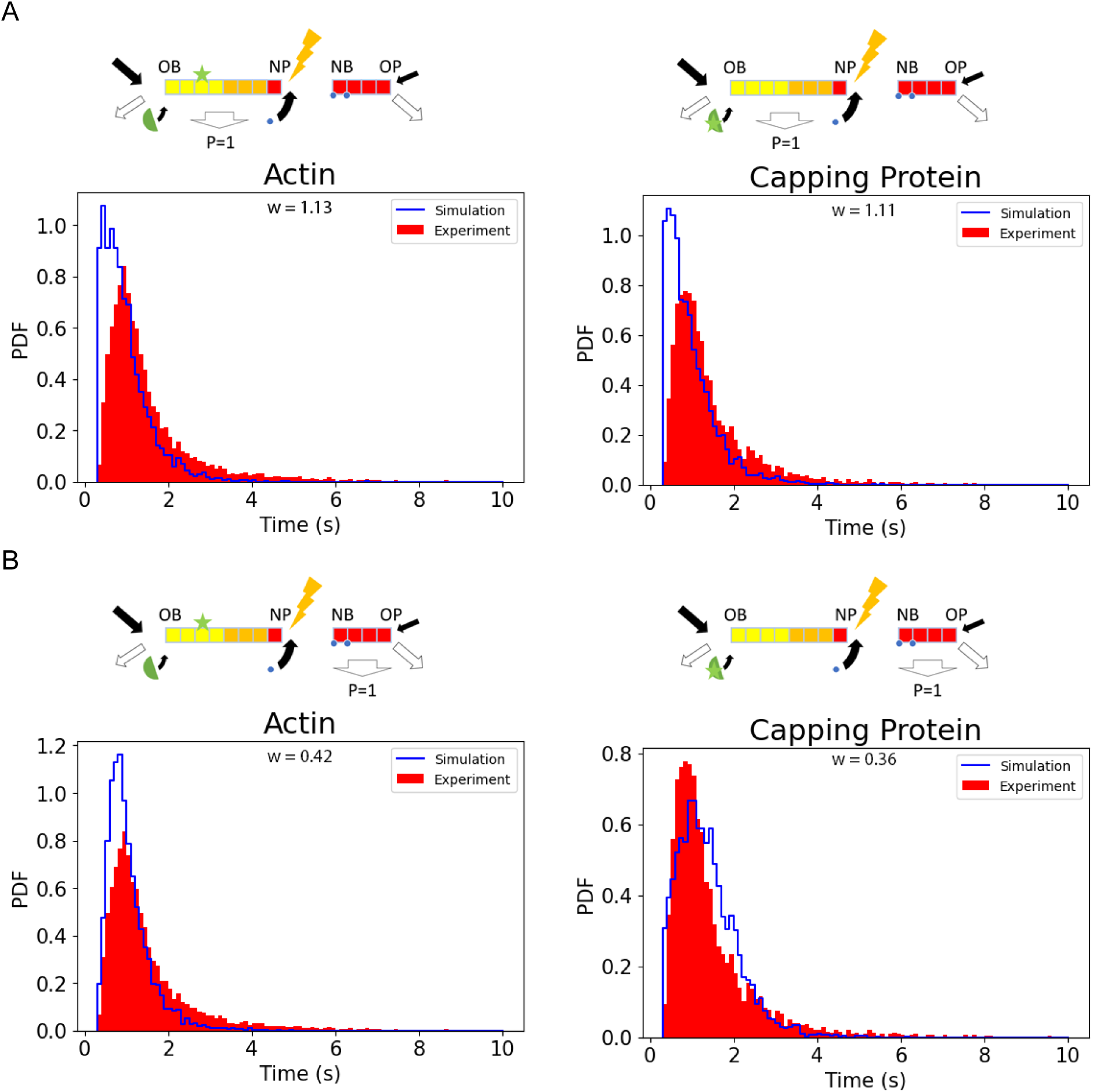
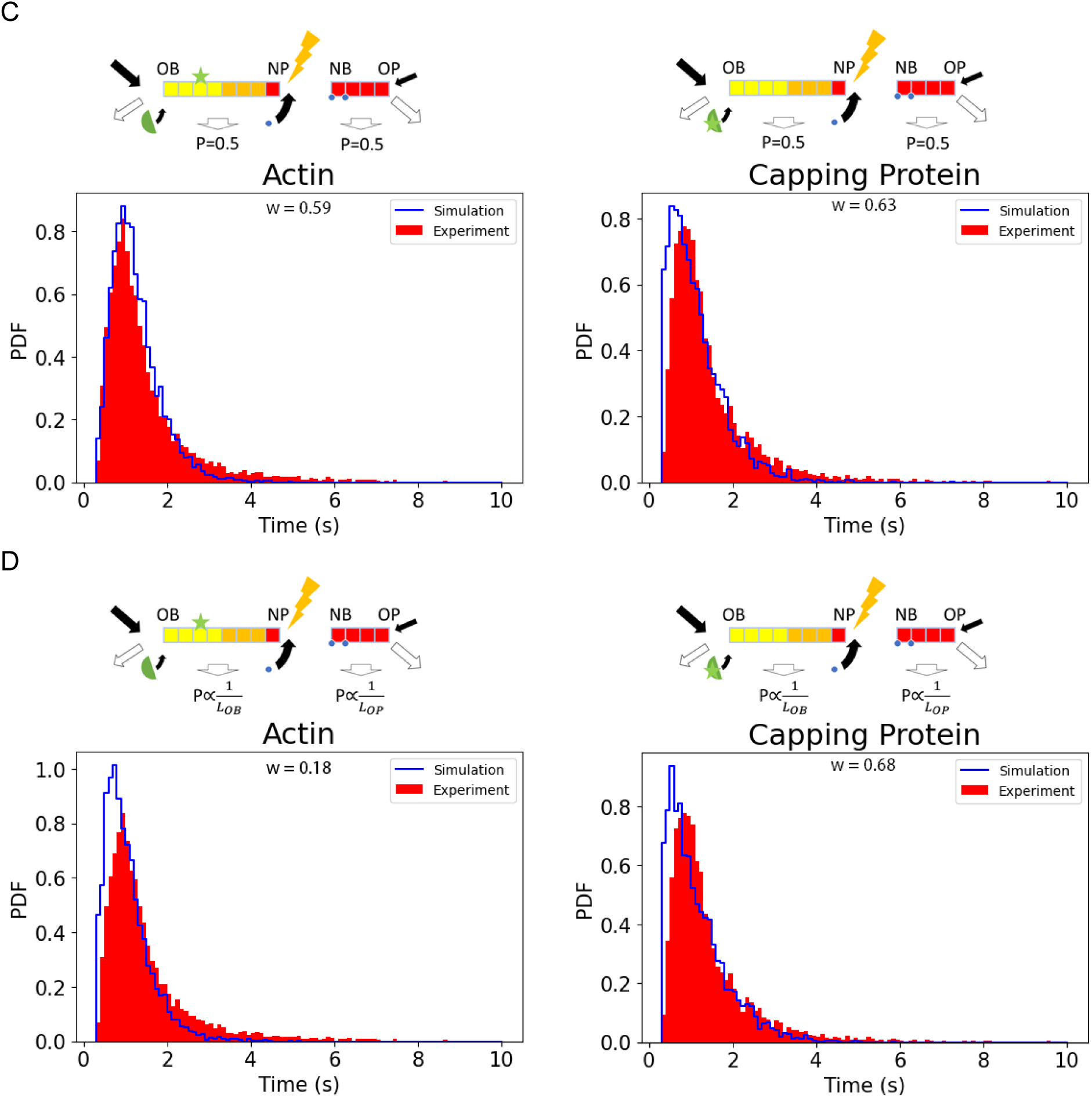
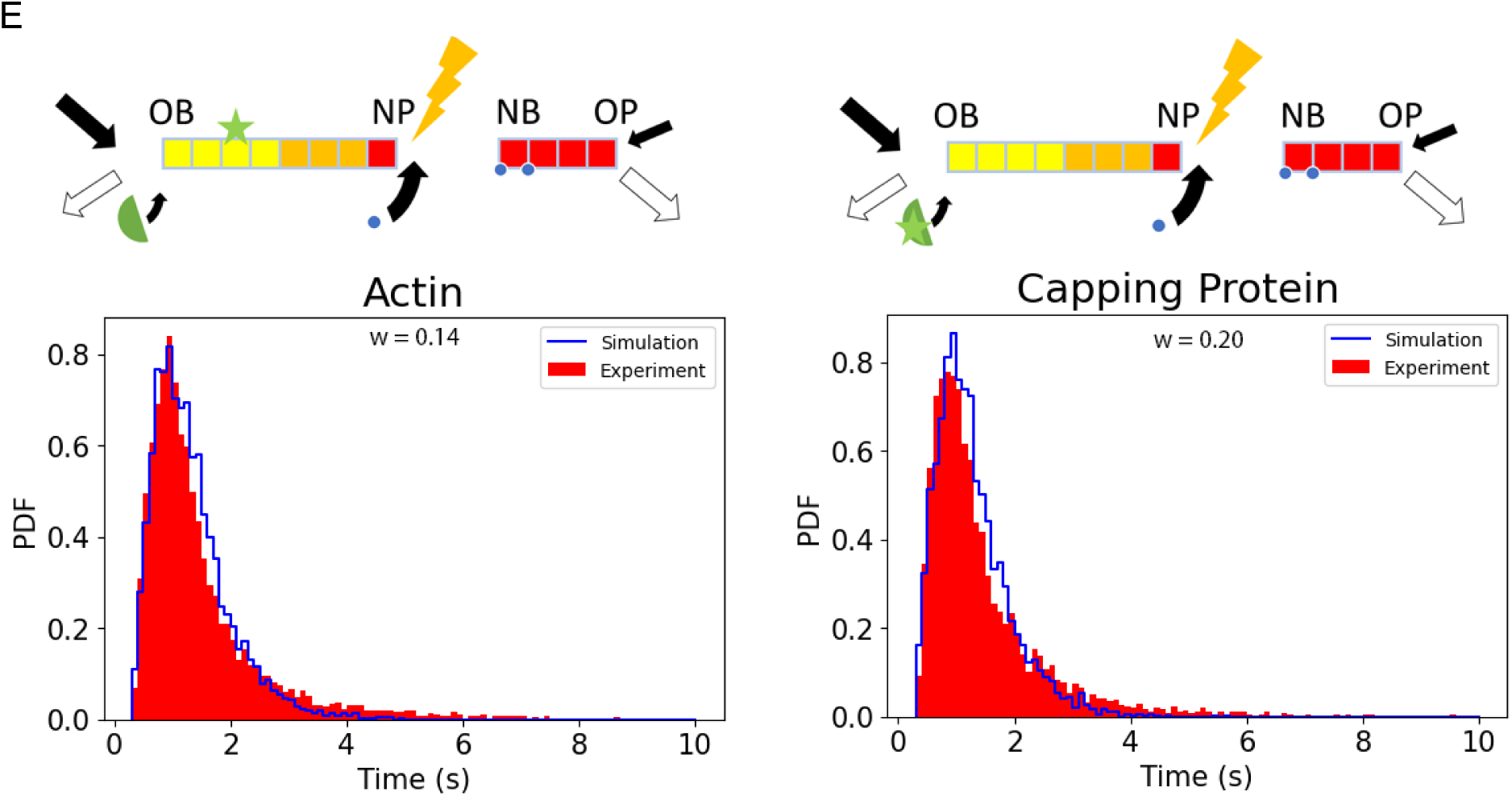
Evaluation of different assumptions on the removal of filament fragments after severing on the dwell time distributions of actin subunits (Left) and the capping protein (Right) A) The fragment that contains the old-barbed end is always discarded; *w*=1.13 for actin and 1.11 for capping protein, B) The fragment that contains the old-pointed end is always discarded; *w*=0.42 for actin and 0.36 for capping protein, C) Either the fragment that contains the old barbed end or the ones that contain the old-pointed end is discarded with equal probability; *w*=0.59 for actin and 0.63 for capping protein, D) Either the fragment that contains the old barbed end or the fragment that contains the pointed end is discarded with a probability inversely proportional to the length of the fragment (i.e. in the depicted schematic, *P*_*OB*_ is smaller than *P*_*OP*_ as the old barbed end fragment is larger than old-pointed end fragment); *w*=0.18 for actin and 0.68 for capping protein. E) For comparison, we have included the cases where we keep both ends of the filament after severing, *w*=0.14 for actin and 0.19 for capping protein (Fig. 2B and Fig. 3A).

**Figure S7.**
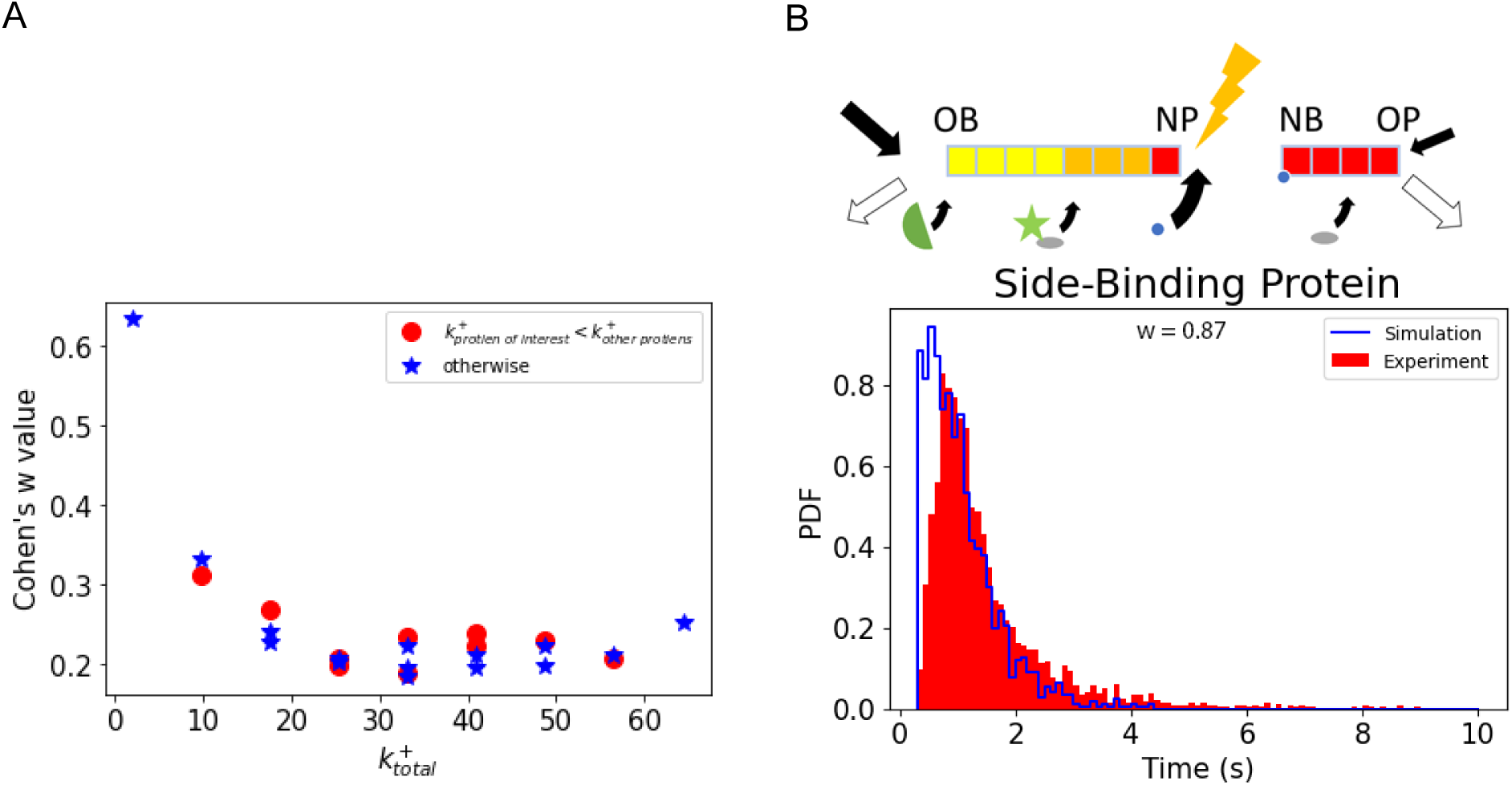
Competition to decorate the actin filaments. A) The competition for decorating a vacant actin subunit is not necessarily between different populations of proteins, since it can be between within the same 0type of protein (e.g., myosin). To demonstrate this, we plot *w* vs 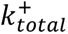, the sum of the association rate of the protein of interest and the association rates of other proteins. As expected, *w* is determined by 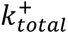 and not just by one of the association rates. To further evaluate this, we have split the data points into two groups. We showed *w* for the cases when the association rate of the protein of interest is lower than the association rate of the other proteins with the red circles, otherwise, we used a blue star. For a specific value of 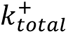 the differences between the two *w* values are less than 0.1. This difference is due to the limited number of data points. The plot has a minimum, which puts an upper limit for 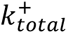. In other words, there should be a small delay between the polymerization of a new actin subunit and the association of the side-binding proteins. B) Dwell time distribution of the side-binding proteins when the competition for the other actin-binding proteins is low. The increased vacant time of actin subunits leads to a shorter average dwell time of the side-binding proteins, *w* = 0.87.

**Figure S8.**
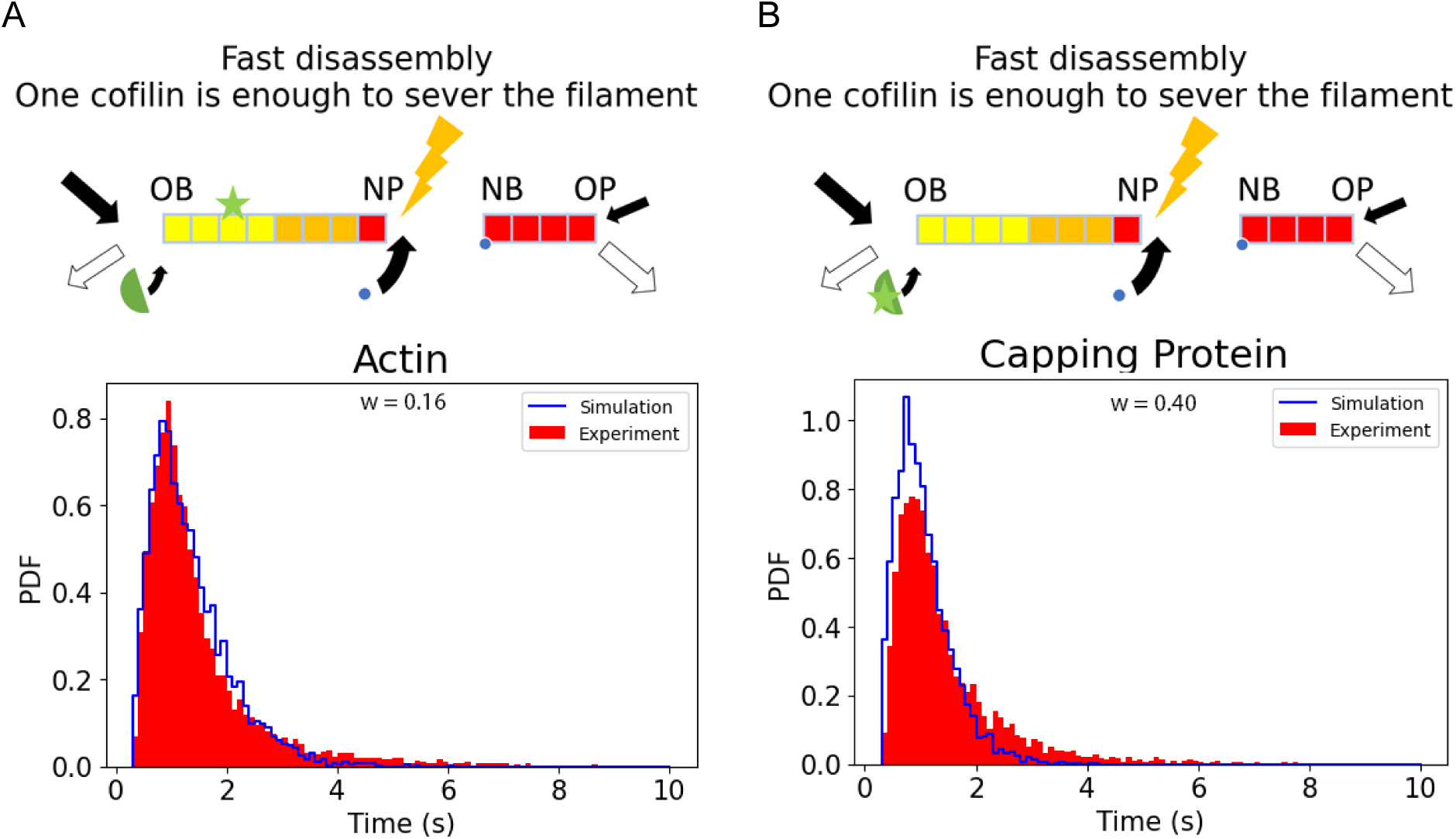
More than one ADF/cofilin-bound actin is required to sever the filament. We ran simulations under the assumption a single cofilin is sufficient to sever a filament (instead of two consecutive ones in the rest of the paper). While the actin subunit dwell time distribution remains similar to experimental data (A), the capping protein dwell time distribution was significantly worse (*w* = 0.40) (B). This result suggests that only one ADF/cofilin-bound actin is not enough to sever the filament.

**Figure S9.**
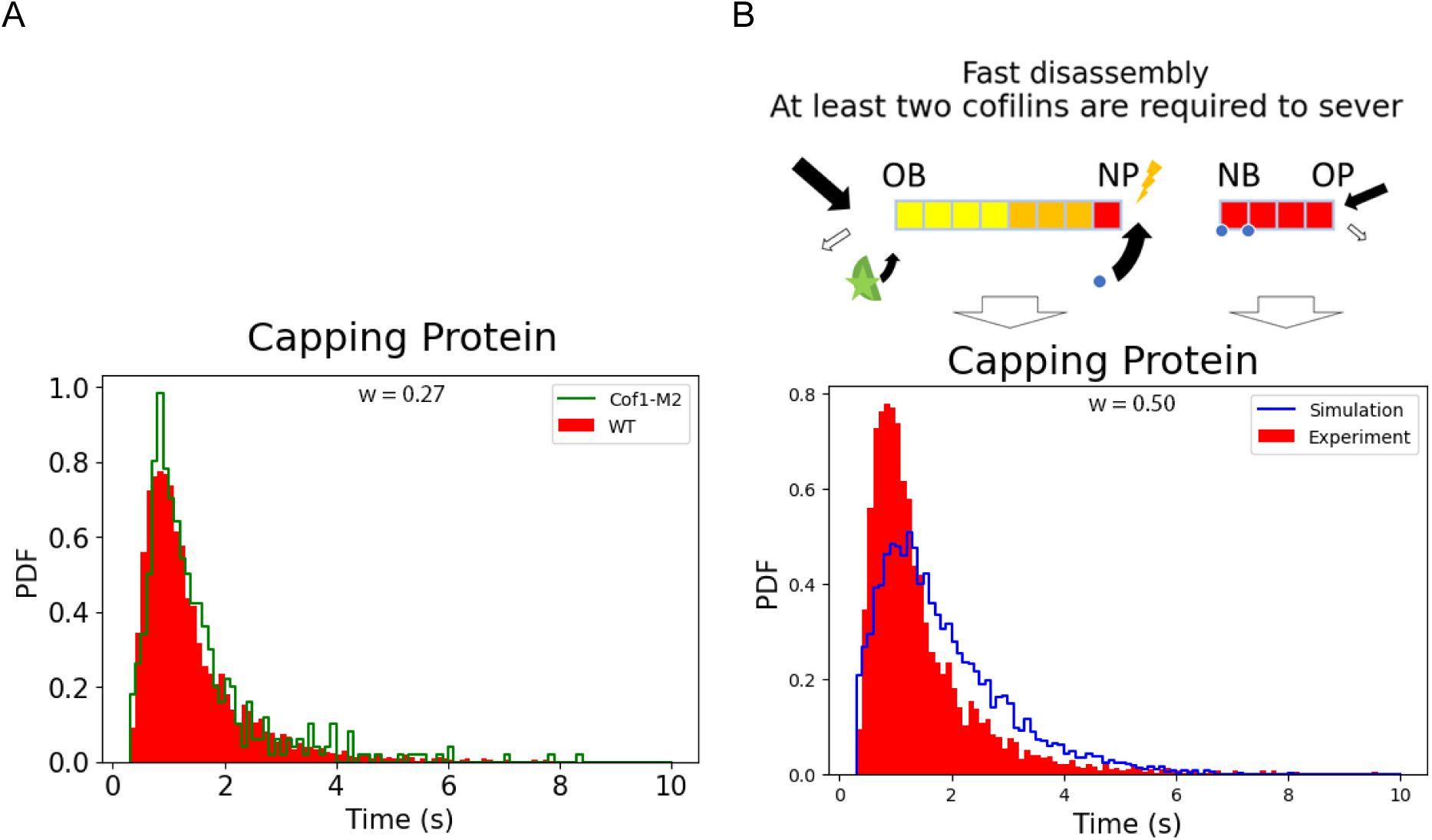
Experimental dwell time distribution for the acting protein subunit Acp1p in the cofilin mutant Cof1-M2. A) While Cof1-M2 severs actin filaments 3.5 times less than wild type *in vitro* (the maximum severing rate is 9×10^-6^s^-1^ versus 32 ×10^-6^s^-1^ (Chen & Pollard, 2011)), it had a small effect on the dwell time of the actin capping protein subunit Acp1p. B) Decreasing the severing rate in the model increased the average capping proteins’ dwell time to 1.7 ± 1.1 s (*w* = 0.5). These data suggest that the *in vivo* severing rate of actin filaments is largely enhanced by other factors (e.g., Aip1, Twinfillin, Srv2/CAP and mechanical forces).

**Figure S10.**
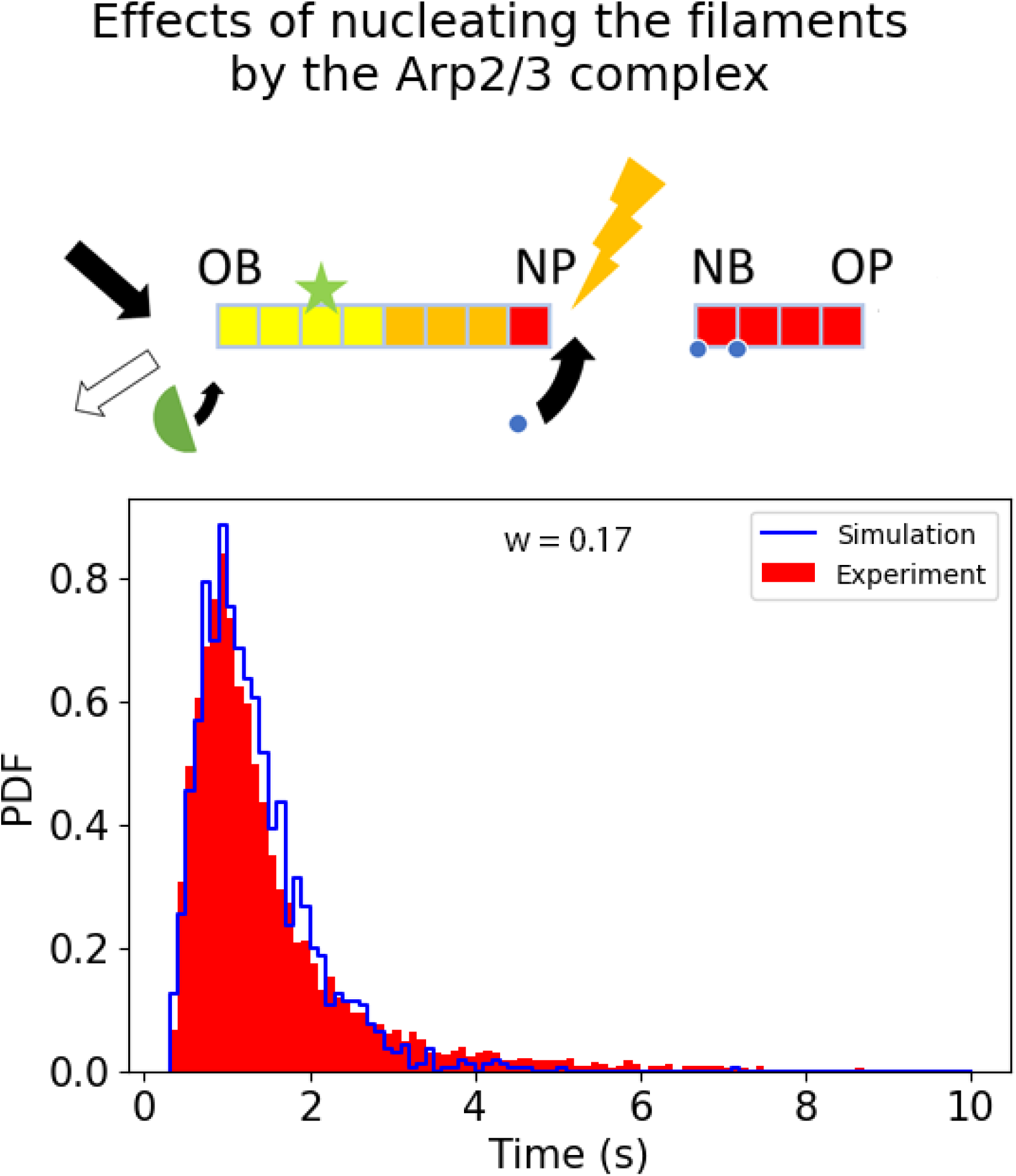
The Arp2/3 complex has a negligible effect on the simulation results. The nucleation of a daughter actin filament by an Arp2/3 complex cap its pointed end and prevents it from depolymerizing. In the extreme case where the Arp2/3 complex does not disassemble from the filament, it stabilizes the filament’s pointed end does. To test if this capping affects the overall actin subunit’s dwell time distribution, we considered the extreme case, where all the pointed ends are stabilized by an Arp2/3 complex. Even in this extreme situation, we obtained a similar distribution as when we did not include the Arp2/3 (*w* = 0.17) by slightly increasing the disassembly rates 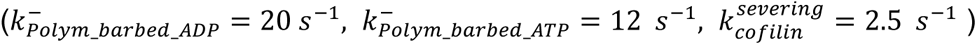.

## Notes

### Competing Interest Statement

The authors have declared no competing interest.

### Summary of Updates

Clarified the text and added new supplemental figures

